# Homozygous haplotype deficiency in Manech Tête Rousse dairy sheep revealed a nonsense variant in *MMUT* gene affecting newborn lamb viability

**DOI:** 10.1101/2023.03.10.531894

**Authors:** Maxime Ben Braiek, Carole Moreno-Romieux, Céline André, Jean-Michel Astruc, Philippe Bardou, Arnaud Bordes, Frédéric Debat, Francis Fidelle, Itsasne Granado–Tajada, Chris Hozé, Florence Plisson-Petit, François Rivemale, Julien Sarry, Némuel Tadi, Florent Woloszyn, Stéphane Fabre

## Abstract

Recessive deleterious variants are known to segregate in livestock populations as in human, and some may cause lethality when homozygous. By scanning the genome of 6,845 Manech Tête Rousse dairy sheep using phased 50k SNP genotypes and pedigree data, we searched for deficiency in homozygous haplotype (DHH). Five Manech Tête Rousse deficient homozygous haplotypes (MTRDHH1 to 5) were identified with a homozygous deficiency ranging from 84% to 100%. These haplotypes are located on OAR1 (MTRDHH2 and 3), OAR10 (MTRDHH4), OAR13 (MTRDHH5) and OAR20 (MTRDHH1), and have frequencies ranging from 7.8% to 16.6%. When comparing at-risk mating between DHH carriers to safe mating between non-carriers, two DHH (MTRDHH1 and 2) showed significant effects on decreasing artificial insemination success and/or increasing stillbirth rate. We particularly investigated the MTRDHH1 haplotype highly increasing stillbirth rate, and we identified a single nucleotide variant (SNV) inducing a premature stop codon (p.Gln409*) in the *MMUT* gene (*methylmalonyl-CoA mutase*) by using a whole genome sequencing (WGS) approach. We generated homozygous lambs for the *MMUT* mutation by oriented mating, and most of them died within the first 24h after birth without any obvious clinical defect. RT-qPCR and western blotting performed on post-mortem liver and kidney biological samples showed a decreased expression of *MMUT* mRNA in the liver and absence of a full-length MMUT protein in mutated homozygous lambs. In parallel, MTRDHH4 and MTRDHH5 showed partial association with variants in *RXFP2* and *ASIP* genes, respectively, already known to control horned/polled and coat color phenotypes in sheep, two morphological traits accounting in the MTR breed standard. Further investigations are needed to identified the supposed recessive deleterious variant hosted by MTRDHH2 and MTRDHH3. Anyway, an appropriate management of these haplotypes/variants in the MTR dairy sheep selection program should increase the overall fertility and lamb survival.

**Author Summary:** In this article, we used reverse genetics screen in ovine using large genotype data available in the framework of genomic selection program in Manech Tête Rousse dairy sheep. We identified five genomic regions with a highly significant deficit in homozygous animal. These regions are thus supposed to host recessive deleterious mutations. In one of these genomic regions, we identified a nonsense mutation in *MMUT* that alters the functioning of this essential gene of cell metabolism, causing perinatal mortality of homozygous lambs. In this work, we also identified other regions possibly associated with morphological appearance part of the breed standard such as polledness and coat color. Increasing knowledge in these genomic regions will help the future genetic management of the Manech Tête Rousse breed, particularly to reduce lamb mortality.

## Introduction

In livestock, genetic selection has largely improved production traits over the past decades, but the last one has seen the emergence of new technological tools allowing to implement the genomic selection that further enhanced the genetic progress [1]. The availability of high-density single nucleotide polymorphism (SNP) chip and the improvement of knowledge of genomes (genome assembly and gene annotations) have allowed to fine-map genomic regions and identify causal variants associated with production traits [1–4]. Despite a successful selection on these traits, undesired decline of fertility was observed [5]. Although the environment explains a large part of performance in ruminants ), genetic studies have made it possible to correct this trend and improve fertility while its heritability is less than 0.05 [6–8].

These studies have shown Mendelian monogenic disorders as one of the causes of fertility failure.

Nowadays two main approaches were broadly developed to identify recessive deleterious variants. The first approach is a “top-down” strategy based on case-control analysis performing genome-wide association [9] when biological samples from affected animals are available. In this method, distinctive phenotypes between non-affected and affected animals are essential. Subsequently, homozygosity mapping approach could be performed to detect homozygous regions in affected animals supposed to host the causal variant, further determined by whole genome sequencing (WGS) data [10,11]. In livestock, Charlier et al. [10] have used this approach for the first time and successfully detected three causal variants in cattle breeds located in *ATP2A1*, *SLC6A5* and *ABCA12* genes responsible for “congenital muscular dystony” types 1 (OMIA 001450-9913) and 2 (OMIA 001451-9913) and “ichthyosis fetalis” (OMIA 002238-9913), respectively. However, this approach showed some limits when biological samples and descriptive phenotypes are not available. To raise this drawback, a second approach called “bottom-up” or reverse genetic screen strategy was developed to specifically identify recessive lethal variants. This strategy, initially developed by VanRaden et al. [12], is based on the exploitation of large number of genotyped animals easily available from genomic selection datasets to detect haplotypes showing deficit in homozygous animals, with a significant deviation from the Hardy-Weinberg equilibrium. Initially, this method was developed to detect embryonic lethal variant but the generalization of the method can also fine-map deleterious variants leading to neonatal or juvenile lethality and morphological disorders. This reverse genetic screen has successfully identified numerous deficient homozygous haplotypes in several species, cattle [12–27], pigs [28,29], chicken [30], turkey [31] and horses [32], and the whole genome sequencing that followed has revealed the associated causative variants hosted by these regions [13,14,18,19,21,24–27,33–41]. We recently, and for the first time, validated this approach in sheep with the identification of 8 independent deficient homozygous haplotypes in the Lacaune dairy breed [42]. Thereafter, focusing on the Lacaune Deficient Homozygous Haplotype 6 (LDHH6, OMIA 002342-9940), we identified a nonsense variant in *CCDC65* gene causing juvenile mortality associated with respiratory distress when homozygous [43].

In the present work, still leveraging genomic selection data, we searched for lethal variants using a reverse genetic screen in Manech Tête Rousse (MTR) dairy sheep. This breed is raised in the French Basque Country and represents the second most important breed by its population size (∼450,000 ewes) in France [44]. Genetic variability is well controlled in MTR breed with an increase by +0.4% of inbreeding per generation over 1999-2009. The effective population size ranges from 110 to 200 according to the estimation methods [44–46]. As for other dairy sheep breeds in France, genomic selection in MTR was implemented in 2017 [47]. The aim of this study was to identify deficient homozygous haplotypes by reverse genetic screen using large amount of genotyping data available in MTR dairy sheep, to test the hypothesis of negative impacts on fertility traits in at-risk matings, to propose relevant candidate genes located in these regions that could host recessive deleterious variants. We particularly focus on one region to identify the associated causal variant from WGS data and manage at-risk mating between carriers to determine the associated phenotype.

## Results

### Identification of deficient homozygous haplotypes in Manech Tête Rousse dairy sheep

Using a reverse genetic screen strategy based on 5,271 genotyped animals belonging to trios, we have detected 150 highly significant Homozygous Haplotype Deficiency (HHD) of 20 SNP markers (listed in S1A Table). These 150 HHD were clustered to five independent regions called “Manech Tête Rousse Deficient Homozygous Haplotype” (MTRDHH). Three haplotypes showed a total deficit in homozygous animals (MTRDHH1, 2 and 3), whereas two haplotypes, MTRDHH4 and 5, only showed a partial deficit (84% and 91%) with 1 and 8 homozygous animals genotyped while 11 and 49 were expected, respectively (Table 1, Fig 1, S1A Table). The complete description of MTRDHH SNP markers (SNP name, SNP allele and position on sheep reference genomes Oar_v3.1, Oar_Rambouillet_v1.0 and ARS-UI_Ramb_v2.0) is available in S1B Table. The different MTRDHH were located on OAR20 (MTRDHH1), OAR1 (MTRDHH2 and 3), OAR10 (MTRDHH4) and OAR13 (MTRDHH5), and their length ranged from 1.1 to 4.6 Mb on Oar_rambouillet_v1.0. The observed frequencies of heterozygous carriers were between 7.8% and 16.6%. MTRDHH2 and MTRDHH3 both located on OAR1 were not in linkage disequilibrium. Consequently, the five MTRDHH identified are likely to harbor five independent variants causal of the observed homozygous deficiency.

**Fig 1.**
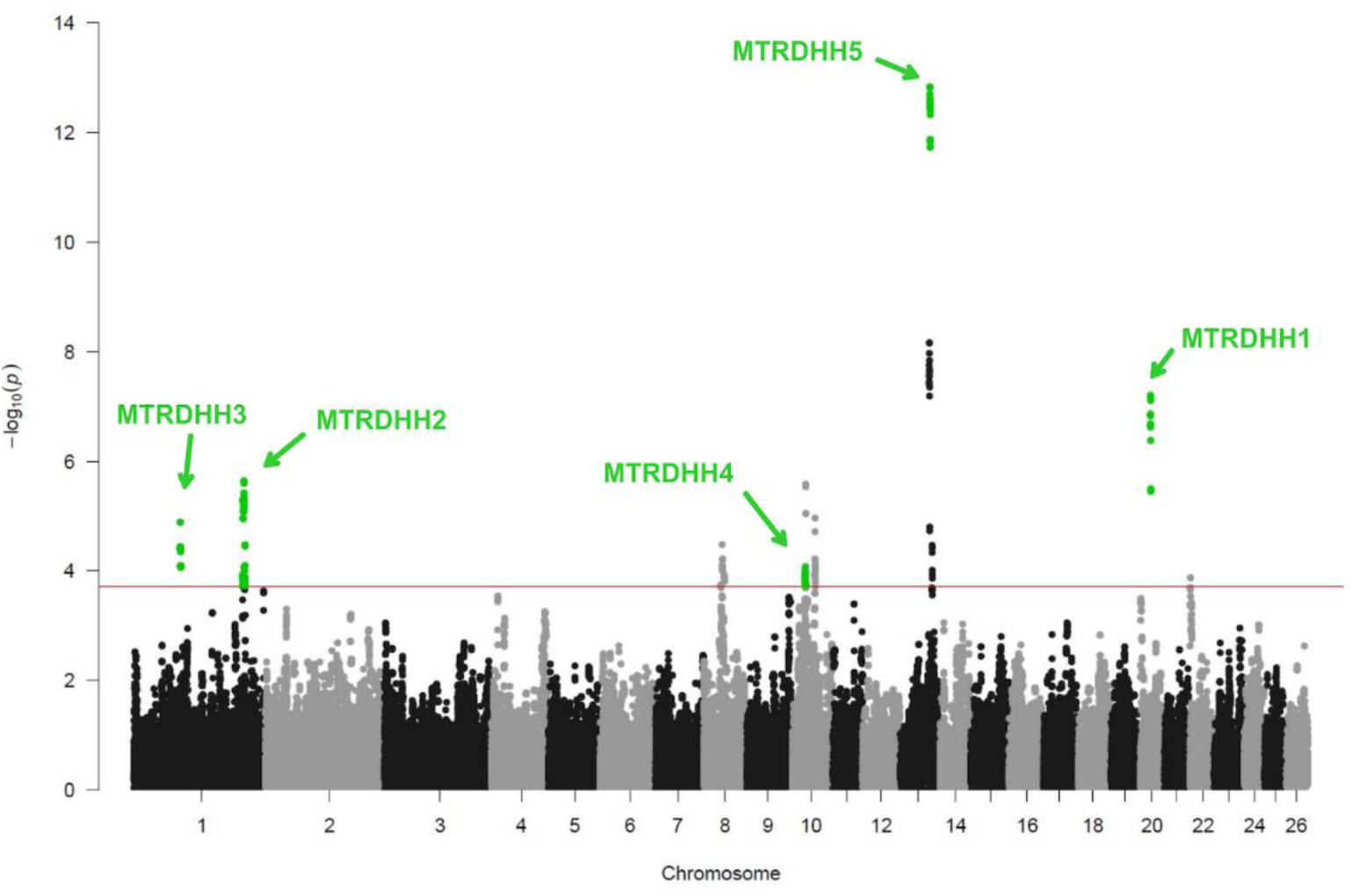
Manhattan plot of HHD identified in Manech Tête Rousse dairy sheep. Each point represents one haplotype of 20 markers with a frequency > 1% in the maternal phase. The red line represents the P-value threshold (1.9 × 10^−4^) used to consider a haplotype with a significant deficit in homozygotes. Only HHD with a deficit in homozygotes ≥ 75% (green dots) were selected and resulted in the identification of 150 significant HHD clustered in 5 regions (MTRDHH1 to 5). Genomic coordinates refer to the sheep reference genome Oar_v3.1.

**Table 1.** List of Manech Tête Rousse deficient homozygous haplotypes.

| Haplotype | OAR | Number of markers | Position (Mb) | Heterozygous carrier frequency (%) | Number of homozygotes |  |  |  |
| --- | --- | --- | --- | --- | --- | --- | --- | --- |
|  |  |  |  |  | <sup>d</sup>Exp | <sup>e</sup>Obs | Deficit (%) | Poisson P-value |
| MTRDHH1 | 20 | 32 | 23.0-25.0 | 9.7 | 13 | 0 | 100 | $2.9 \times 10^{-6}$ |
| MTRDHH2 | 1 | 66 | 251.9-256.4 | 8.7 | 10 | 0 | 100 | $3.8 \times 10^{-5}$ |
| MTRDHH3 | 1 | 39 | 103.8-106.6 | 7.8 | 9 | 0 | 100 | $9.6 \times 10^{-5}$ |
| MTRDHH4 | 10 | 26 | 30.5-31.5 | 8.7 | 11 | 1 | 91 | $1.5 \times 10^{-4}$ |
| MTRDHH5 | 13 | 29 | 64.3-67.2 | 16.6 | 49 | 8 | 84 | $5.3 \times 10^{-13}$ |

<sup>a</sup>MTRDHH haplotypes and SNP markers composition are details in S1 Table
<sup>b</sup>Position on ovine genome assembly Oar\_rambouillet\_v1.0
<sup>c</sup>Frequency of carriers in the entire genotyping population (n=6,845)
<sup>d</sup>Expected
<sup>e</sup>Observed

### Impact of MTRDHH on fertility traits

In order to identify a putative lethal effect of the five MTRDHH, two recorded fertility traits were analyzed: artificial insemination success, a proxy for embryonic loss (AIS: 330,844 matings) and stillbirth rate associated with perinatal lethality (SBR: 201,637 matings) (Fig 2). The average AIS of the population was 60.9%. When comparing at-risk and safe matings, only MTRDHH2 showed a significant decrease of -3.3% of AIS (P=3.5×10^-4^) in at-risk matings. The average SBR of the population was 7.5%. As described in Fig 2, MTRDHH1 and 2 showed a huge increase in SBR with +7.5% (P=4.0×10^-24^) and +4.3% (P=1.3×10^-6^) in at-risk matings compared to safe matings, respectively. The three other haplotypes showed no significant impact on the fertility traits studied.

**Fig 2.**
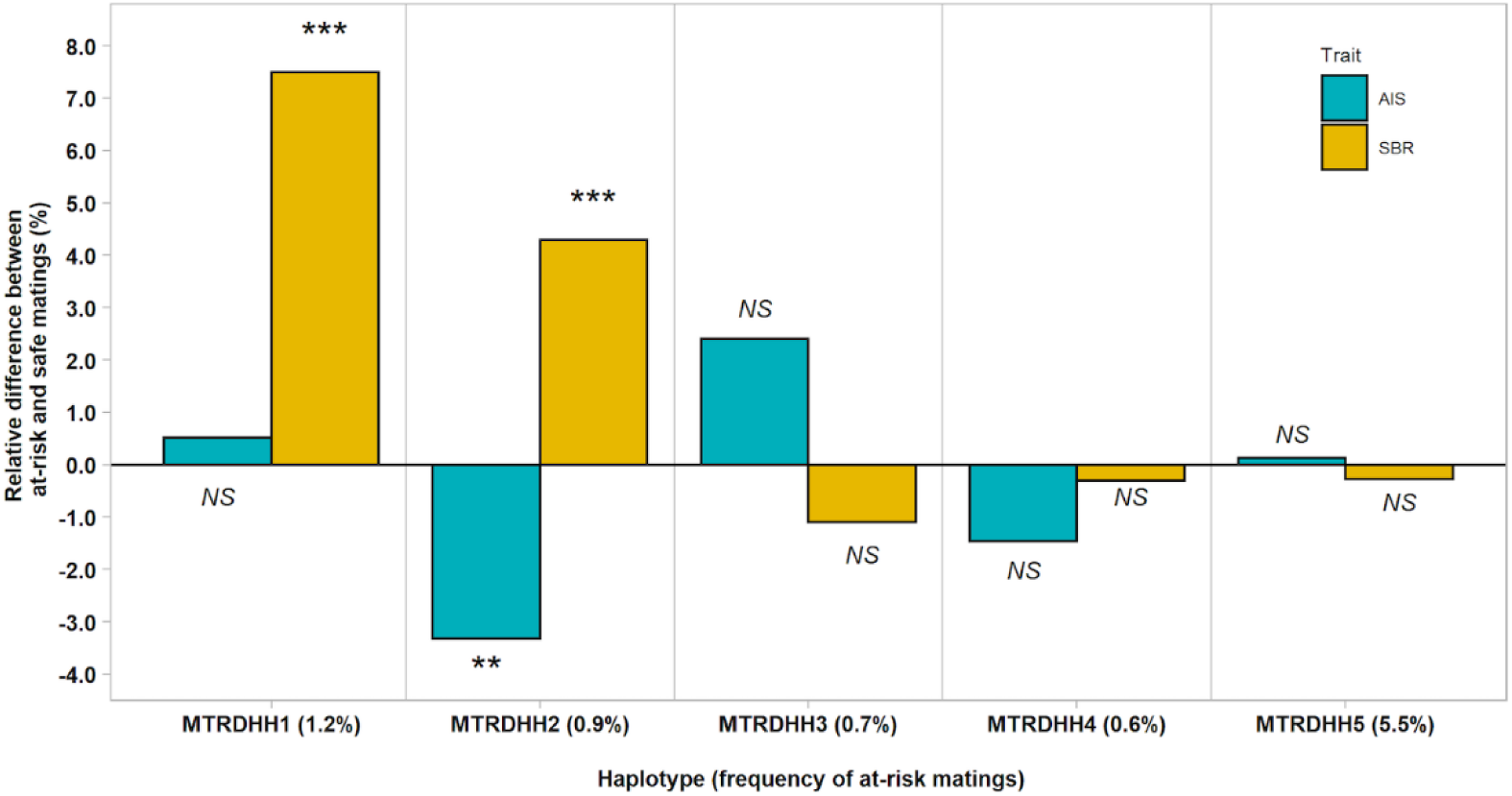
Effects of MTRDHH on artificial insemination success (AIS) and stillbirth rate (SBR) between at-risk and safe matings. For each MTRDHH, the frequency of at-risk matings is shown in parentheses. Significant effects are indicated by the corrected P-value for multiple tests with a threshold set at α=0.1% (**), 0.01% (***). NS, not significant.

### Pleiotropic effects of MTRDHH on milk production traits

Six dairy traits are routinely included in genomic evaluation of the French dairy sheep. Thus, standardized daughter yield deviation (sDYD) of five milk production traits (milk, fat and protein yields, and fat and protein contents) and lactation somatic cell score (a proxy for udder health) were compared between carrier and non-carrier rams for each of the 5 MTRDHH evidenced (Fig 3). Among the five haplotypes, three were associated with significant effect on sDYD. Daughters of MTRDHH2 carrier rams showed a significant increase in milk production (sDYD +0.06, P=6.5×10^-3^) but a decrease in protein content (sDYD -0.11, P=4.7×10^-4^). For MTRDHH4, there was a significant increase in lactation somatic cell score (sDYD -0.13, P=2.4×10^-3^), and daughters of MTRDHH5 carrier rams showed higher fat yield (sDYD -0.06, P=9.6×10^-3^). Additionally, the total merit genomic index, named ISOLg, was extracted from each male lamb of the 2021 genomic selection cohort to estimate the MTRDHH impact on the genetic gain of the selected traits. No significant difference was observed on ISOLg between heterozygous carrier and non-carrier lambs for each MTRDHH (S1 Fig).

**Fig 3.**
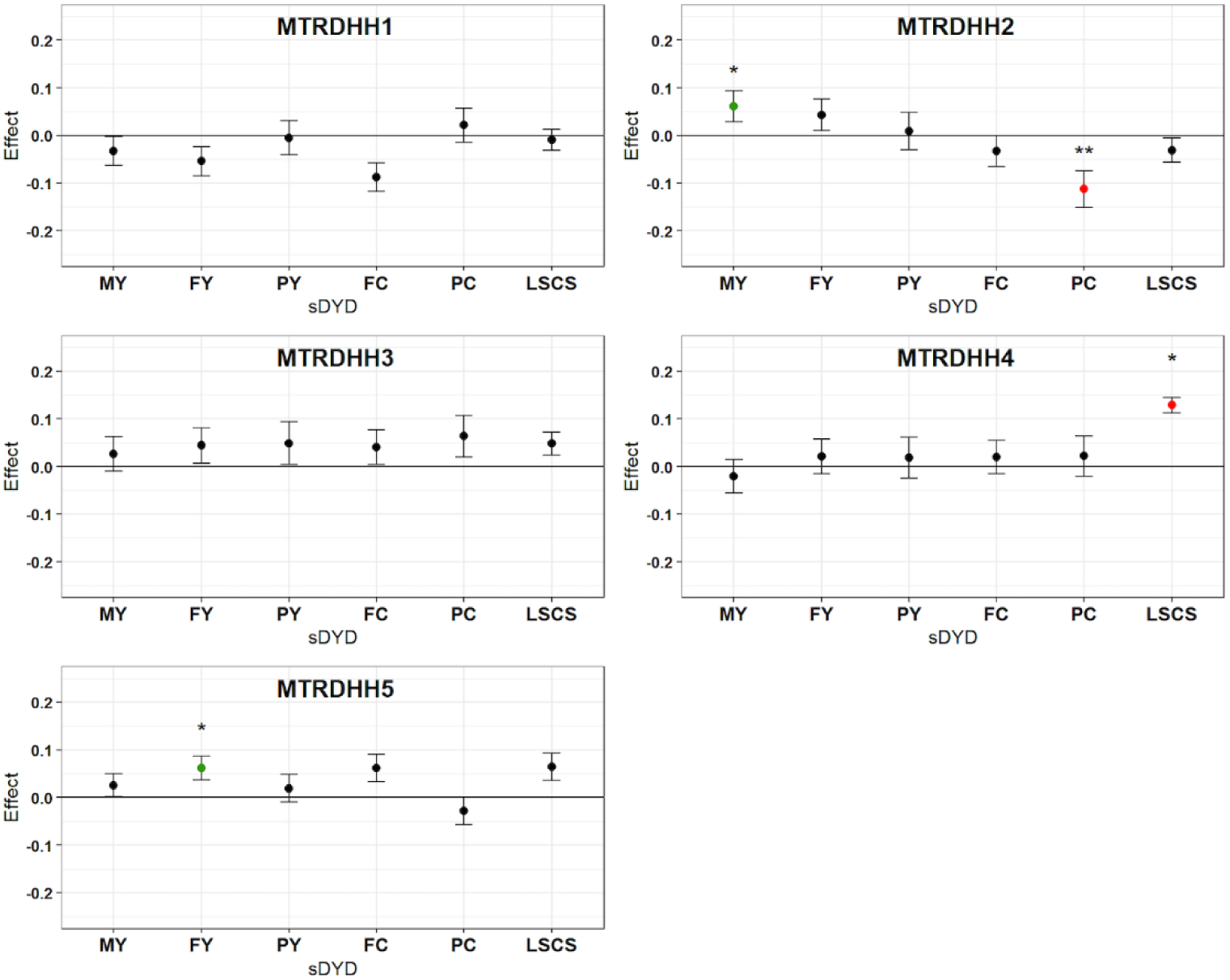
sDYD relative difference between heterozygous and non-carrier rams for 6 traits under selection. MY: milk yield, FY: fat yield, PY: protein yield, FC: fat content, PC: protein content, LSCS: lactation somatic cell score, sDYD standardized daughter yield deviation (DYD divided by genetic standard deviation). sDYD relative difference value is obtained from lsmeans estimate according to mating class. Significant effects are indicated by the corrected P-value for multiple tests with a threshold set at α=5% (*), 0.1% (**). Error bars indicate standard errors. Significant favorable effects of heterozygous are in green while significant unfavorable effects are in red.

### Evolution of the MTRDHH frequencies in the population

Since the implementation of genomic selection in 2017, all candidate rams coming from elite mating were genotyped on low density SNP chip at 7 days old, representating the genetic diversity disseminated by AI in the selection scheme. As shown in Fig 4, the frequencies of the MTRDHH heterozygous carriers were quite stable during the last five years around 6.8, 7.4, 9.3, 10.2 and 16.1% for MTRDHH1, 2, 3, 4 and 5, respectively. Nevertheless, we can notice a spectacular increase in the frequency of MTRDHH3 from 2.8% to 10.9% when comparing 2017 and 2018.

**Fig 4.**
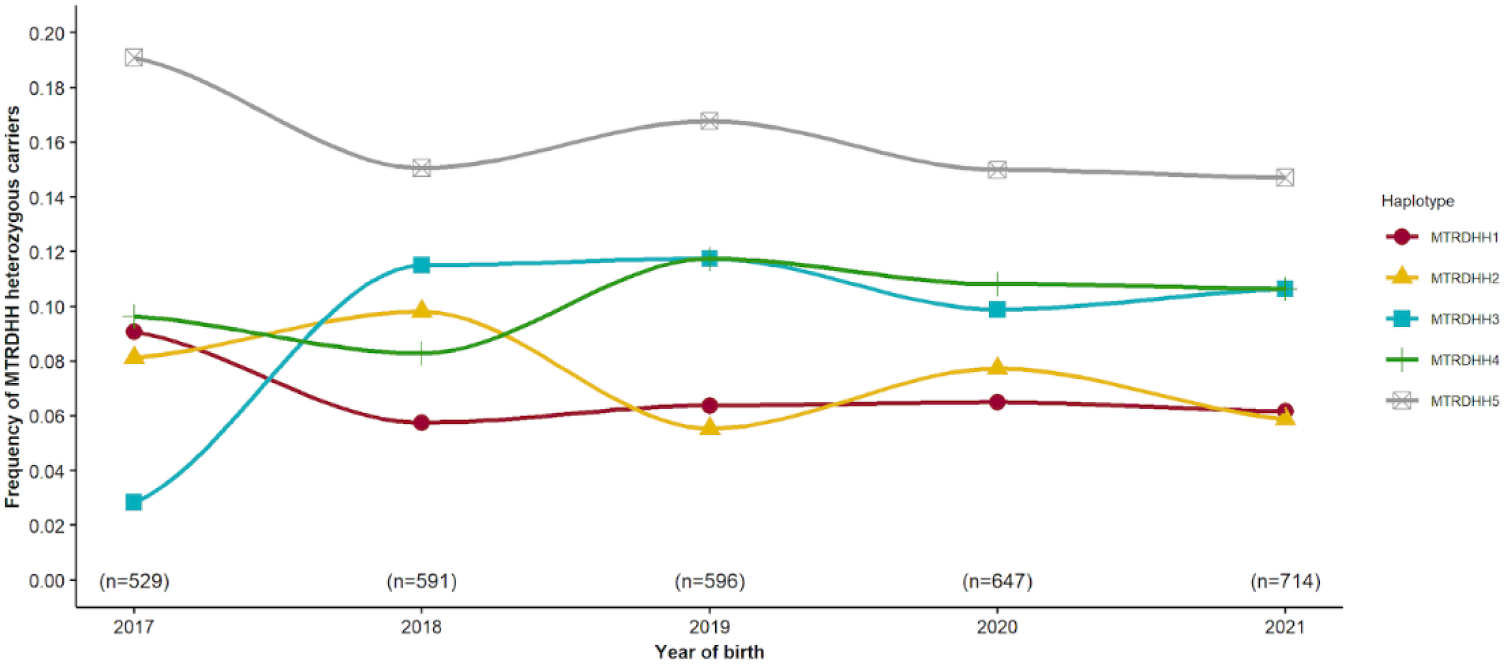
Evolution of the MTRDHH heterozygous carrier frequencies between 2017 and 2021 in Manech Tête Rousse male lambs. For each year the effective refers to all candidates genotyped to enter in the genomic selection scheme.

### Candidate genes located in MTRHH regions

Within the five MTRDHH genomic regions extended by 1Mb on each side, 408 protein coding genes are annotated (S2 Table). When available, information on mouse phenotypes (including lethal phenotypes) and association with mammalian genetic disorders were extracted for each gene using MGI, IMPC, OMIM and OMIA databases. Among the 408 genes, we highlighted 64 genes involved in lethal phenotypes in knock-out mice, and 45 genes associated with human genetic disorders. Twenty-three relevant candidate genes were identified by the intersection of both information (Fig 5). In addition, 7 genes are known to be associated with genetic disorders or morphological traits in livestock (*GJA5*, *ITGA10*, *ADAMTLS4*, *RXFP2*, *KIF3B*, *ASIP* and *CEP250*). Overall, these candidate genes are involved in essential functions such as transcription (*POLR3GL*, *SF3B4*, *PRPF3*, *ASXL1* and *DNMT3B*), cell division (*POGZ*, *BRCA2*, *PIGU* and *CEP250*), basal metabolic processes (*MMUT*, *SLC33A1*, *TARS2* and *AHCY*), cell structure and signaling (*CD2AP*, *PKHD1*, *GJA5*, *ITGA10*, *ECM1*, *GJA5*, *ITGA10*, *ECM1*, *PRUNE1*, *RXFP2*, *KL*, *POFUT1*, *KIF3B*, *ASIP* and *GSS*), and DNA/protein binding (*TFAP2B* and *PEX11B*).

**Fig 5.**
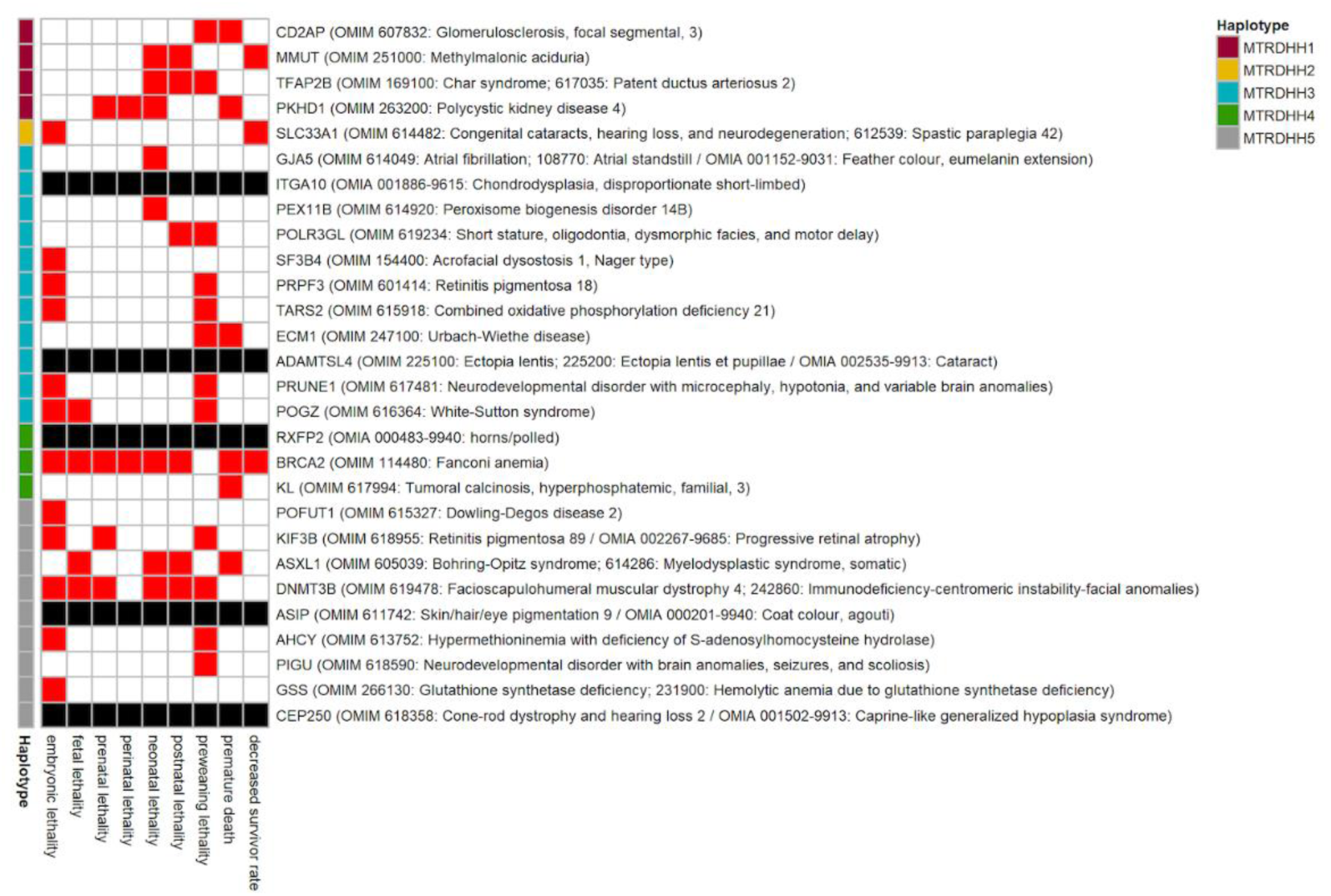
Relevant candidate genes located in MTRDHH implicated in lethal mouse phenotype and/or associated with mammalian genetic disorders. Red boxes indicate time of known lethality stages in mouse when the gene is knocked-out from MGI/IMPC databases. When the gene is associated with mammalian genetic disorders, the OMIM and/or OMIA trait phenotypes are described in parentheses. Black boxes indicate a gene implicated in animal genetic disorder (OMIA) but with no lethal phenotype observed in mouse.

### Known variants in *RXFP2* and *ASIP* genes are partially associated with MTRDHH4 and MTRDHH5

The above list of candidate genes particularly highlighted *RXFP2* (within MTRDHH4) and *ASIP* (within MTRDHH5) genes well-known to impact sheep morphological traits such as polledness [48] and black coat color [49,50], respectively. Interestingly, horned female and black males are not desired in accordance with the MTR breed standards. Using the 22 WGS of MTR animals, we searched for the 1.8kb insertion in the 3’-UTR of *RXFP2* associated with polledness (OMIA 000483−9940, [48]) and the different variants affecting *ASIP* leading to recessive black coat color (OMIA 000201-9940); OAR13:g.66,475,132_66,475,136del and g.66,474,980T>A, Oar_rambouillet_v1.0) [49,50]. S2 Fig shows the segregation of these variant among the 22 sequenced animals and their relationship with the status of MTRDHH4 and 5, but with no obvious association. This being made on a reduced number of animals, we specifically genotyped a larger set of animals (n=714 male lambs born in 2021) for the 1.8kb insertion in *RXFP2* and the two variants in *ASIP*. Using this cohort, we evidenced a partial association between MTRDHH4 and the 1.8kb insertion in the 3’-UTR of *RXFP2,* none heterozygous MTRDHH4/+ being Del/Del corresponding to the horned phenotype (Fig 6). Concerning *ASIP* variants, MTRDHH5 was also partially associated with the 5pb deletion. Indeed, 89% of the MTRDHH5 heterozygous carriers were heterozygous for the 5pb deletion and none Ins/Ins animal was MTRDHH5 carrier. In contrast, we identified almost the same proportion of genotypes for g.66,474,980T>A in MTRDHH5 carriers and non-carriers (+/+), suggesting that MTRDHH5 was not linked to this polymorphism (Fig 6C).

**Fig 6.**
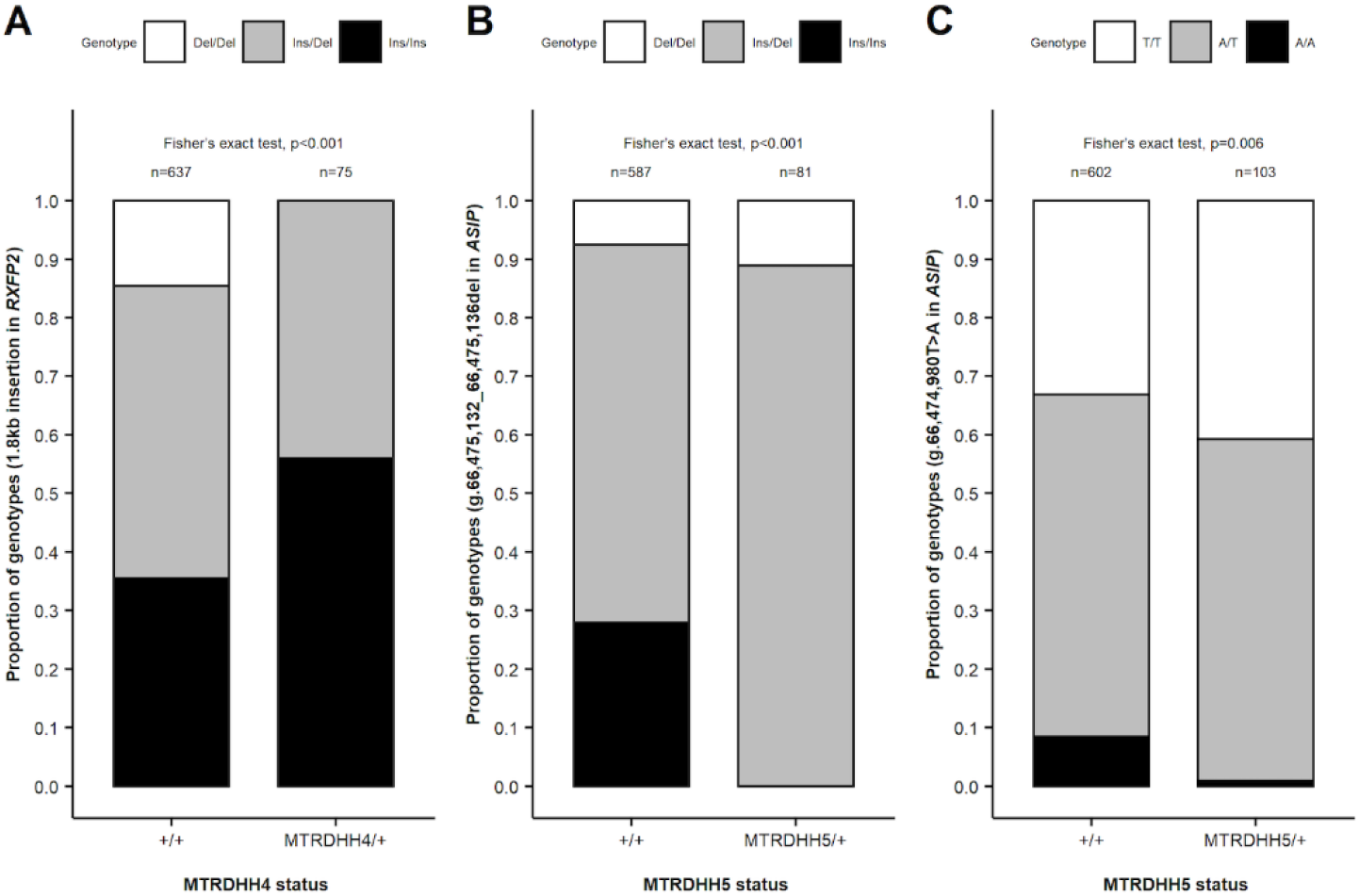
Association between MTRDHH4-5 and causal variants involved in morphological traits. (A) Association between MTRDHH4 status and 1.8kb insertion in the 3’-UTR of *RXFP2* associated with polledness. Association between MTRDHH5 status and (B) OAR13:g.66,475,132_66,475,136del and (C) OAR13:g.66,474,980T>A in *ASIP* associated with black coat color. Coordinates refer to sheep genome Oar_rambouillet_v1.0.

### Identification of a nonsense variant in *MMUT* gene associated with MTRDHH1

With the most important impact on stillbirth rate increased by +7.5% in at-risk matings (Fig 2), we particularly focused on MTRDHH1 that may represent a putative recessive lethal haplotype. In order to identify the MTRDHH1 causal mutation, we have considered biallelic variants (SNP and InDels) for 100 ovine WGS containing the 22 Manech Tête Rousse dairy sheep, and among them two heterozygous carriers of the MTRDHH1 haplotype. Within the MTRDHH1 region extended by 1Mb on each side, 78,019 variants were called with a quality score >30, call rate >95% and only four candidate variants had a perfect correlation (r^2^=1) between biallelic variant genotypes and MTRDHH1 status (Table 2, Fig 7A). Among those candidate variants, we identified two small insertions, one intergenic single nucleotide variant (SNV), and one nonsense (stop-gain) SNV located in the *Methylmalonyl-CoA Mutase* (*MMUT*) gene. This latter SNV (NC_040271.1: g.23,776,347G>A; XM_004018875.4: c.1225C>T; Fig 7B, C) in *MMUT* is predicted to create a premature stop codon at position 409 encoded by exon 6 (XP_004018923.1:p.Gln409*) whereas the full protein length is composed of 750 amino acids (Fig 7D). The variant would disrupt the methylmalonic coenzyme-A mutase domain and would result in the loss of the vitamin B12 binding domain.

**Fig 7.**
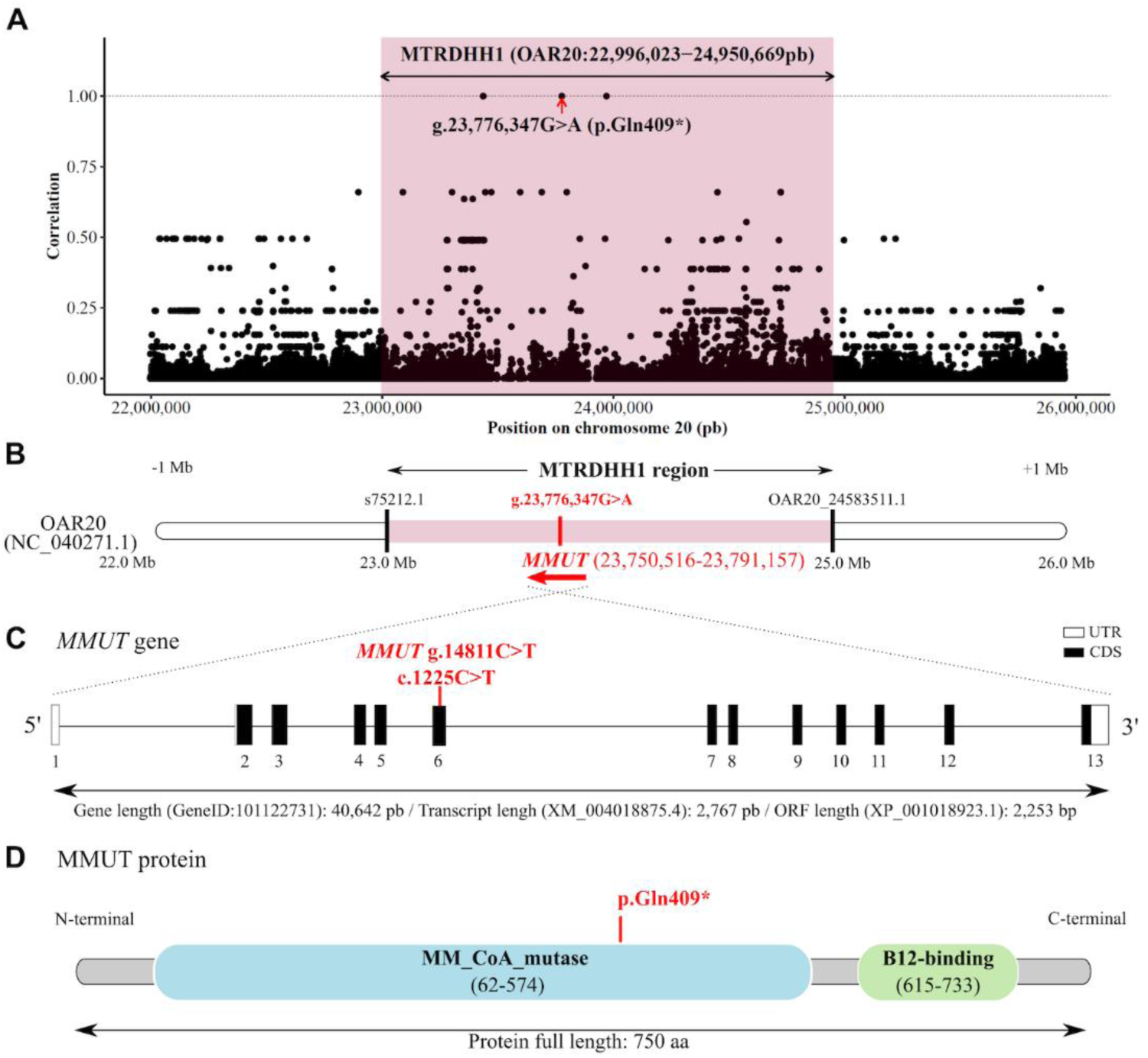
Nonsense variant in *MMUT* gene within MTRDHH1 genomic region. (A) Scatter plot showing the correlation between MTRDHH1 status (NC_040271.1, OAR20:22,996,023-24,950,669 extended from each side by 1 Mb) and genotype of variants from 100 whole genome sequenced animals. Each dot represents one variant. (B) Position of the *MMUT* gene within the MTRDHH1 haplotype. Black bars indicate the first and the last markers of the Illumina Ovine SNP50 BeadChip defining the limits of MTRDHH1 (S1B Table). (C) *MMUT* gene structure (GeneID: 101122731) and localization of the *MMUT* C>T polymorphism identified in the sixth exon (XM_004018875.4). (UTR: untranslated region; CDS: coding sequence) (D) MMUT protein (XP_004018923.1) with Pfam domain annotations (accession number: A0A6P3T7X3_SHEEP) composed of methylmalonyl-CoA mutase (PF01642) and B12-binding (PF02310) domains. The mutation creates a premature stop-gain at amino-acid position 409.

**Table 2.** Candidate variants located in MTRDHH1.

| Position | Ref/Alt | Quality score | Location Annotation | Functional Consequence<sup>a</sup> |
| --- | --- | --- | --- | --- |
| 23,436,234 | G/GTCACA | 385.8 | Intergenic | Modifier |
| 23,436,236 | T/TTTGTG | 385.8 | Intergenic | Modifier |
| 23,776,347 | G/A | 146.0 | Exonic, *MMUT* (c.1225C>T) | High, stop-gain (p.Gln409*) |
| 23,969,676 | C/T | 370.3 | Intergenic | Modifier |

<sup>a</sup>Variant annotation and effect predicted by SnpEff [51].

In order to validate the association between the *MMUT* variant and MTRDHH1, we genotyped the cohort of male lambs born in 2021 (n=714) with a specific genotyping test for the *MMUT* g.23,776,347G>A SNV. The A variant allele frequency was 3.8%. All these animals have a known status at the MTRDHH1 locus and the contingency table indicates a clear association between the MTRDHH1 status and *MMUT* variant genotypes (Fig 8, Fischer’s exact test p<0.0001). However, 15 animals showed discrepancy between *MMUT* and MTRDHH1 genotypes, supposed to be in perfect linkage disequilibrium. A specific focus on haplotypes carried by these animals in the MTRDHH1 region from marker 1 (s75212.1) to marker 32 (OAR20_24583511.1) showed that the 14 animals heterozygous for the variant exhibited shorter recombinant versions of the MTRDHH1 haplotype (S3 Fig). Nonetheless, one animal was heterozygous for MTRDHH1 but did not carry the *MMUT* variant.

**Fig 8.**
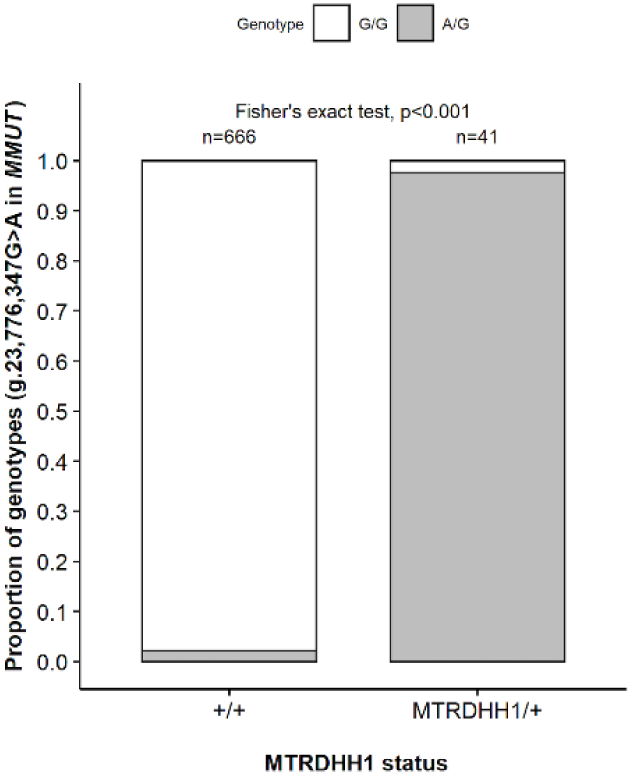
Association of *MMUT* SNV genotypes with MTRDHH1 status. +/+: non-carriers; MTRDHH1/+: heterozygous carriers and MTRDHH1/MTRDHH1: homozygous carriers (Fisher’s exact test, p<0.001, without the homozygous MTRDHH1 carriers).

### Occurrence of the *MMUT* SNV in an ovine diversity panel

An ovine diversity panel composed of 25 French sheep breeds, including MTR, and 3 Latxa Spanish sheep breeds related to MTR was used to genotype the *MMUT* g.23,776,347G>A SNV (Table 3). As expected, some MTR animals (n=5) from this panel were evidenced as heterozygous carriers, and the variant was also detected in one animal of the Spanish Latxa Cara Rubia population. All the other animals tested did not carry the polymorphism.

**Table 3.**
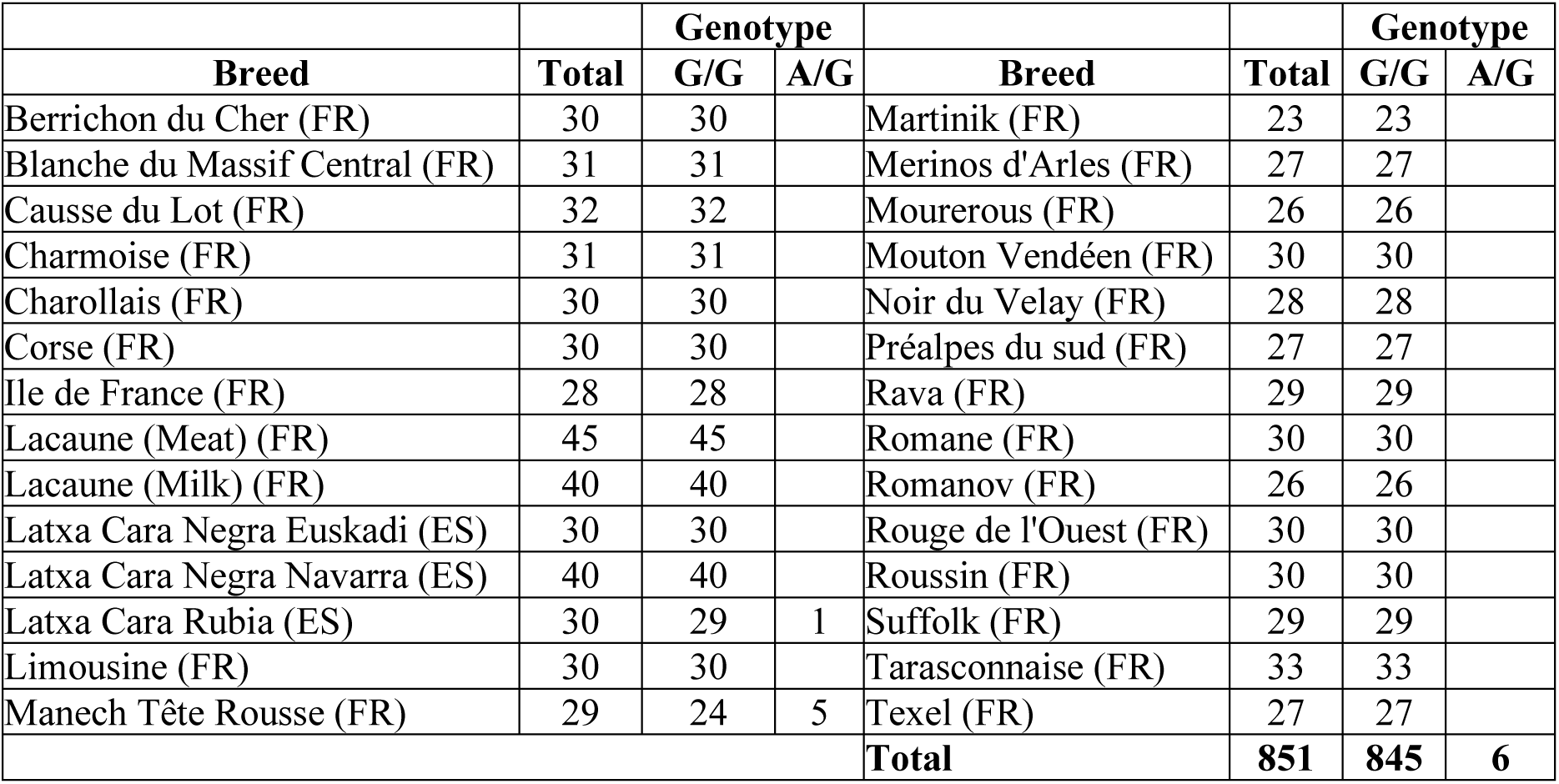
*MMUT* SNV genotype distribution from a DNA diversity panel of French (FR) and Spanish (ES) ovine breeds.

### Viability of homozygous lambs for the *MMUT* variant

To validate the impact of the *MMUT* A variant allele we have manage an oriented mating between heterozygous carriers to generate homozygous lambs. Blood samples were collected from 181 MTR ewes, daughters of MTRDHH1 carrier sires, in 6 private farms. The *MMUT* SNV specific genotyping identified 82 heterozygous ewes. Among these ewes, 73 were raised in the 6 private farms (Experiment 1) and 9 were moved into an INRAE experimental farm (Experiment 2). All ewes were artificially inseminated with fresh semen from *MMUT* g.23776347G/A heterozygous rams. Forty-five days after AI, 44 and 5 were diagnosed as pregnant by ultrasonography in Experiment 1 and 2, respectively. This corresponds to an AIS of 59.8% in accordance with the average AIS of 60.9% determined previously in the whole population. In experiment 1, only 37 among the 44 pregnant ewes were monitored after gestation diagnosis and resulted in the birth of 59 lambs (mean prolificacy of 1.6, litter size ranging from 1 to 3) with a gestation length between 139 and 159 days. In experiment 2, the 5 pregnant ewes gave birth to 13 lambs (mean prolificacy of 2.6, litter size ranging from 2 to 4) with a gestation length between 151 and 157 days. No abortion during the five months of gestation was observed. Finally, 72 lambs (52% males and 48% females) were born and an ear punch was collected for genotyping the *MMUT* SNV (Table 4). The distribution of genotypes did not differ between the two experiments (Fisher’s exact test, p=0.686). In total, 21 lambs were genotyped homozygous carriers (A/A), 29 heterozygous carriers (A/G) and 21 homozygous non-carriers (G/G). All lambs were monitored during the 0-30 days period until weaning. Twenty-five lambs died during this period representing a huge mortality rate of 34.7% (Fig 9). Contingency table between lamb genotypes (A/A, G/A, G/G) and viability (alive or dead) indicated a higher mortality rate for homozygous A/A lambs (Table 4, Fisher’s exact test, p<0.001). Indeed, the A/A dead lambs accounted for 78% of the whole lamb mortality. The death of A/A homozygous lambs occurred very soon after birth within the first 24 hours. Clinical examination of dead lambs did not allow us to identify specific symptoms. Two homozygous A/A lambs have passed the weaning age (around 4 weeks). Additionally, in Experiment 2, the 13 lambs were weighted at birth (males: 4.0 ± 0.9 kg, females: 2.9 ± 1.4 kg) and A/A lambs had significantly lower birth weight compared to the other genotypes, regardless gender (Wilcoxon’s non parametric test, p=0.019) (Fig 10).

**Fig 9.**
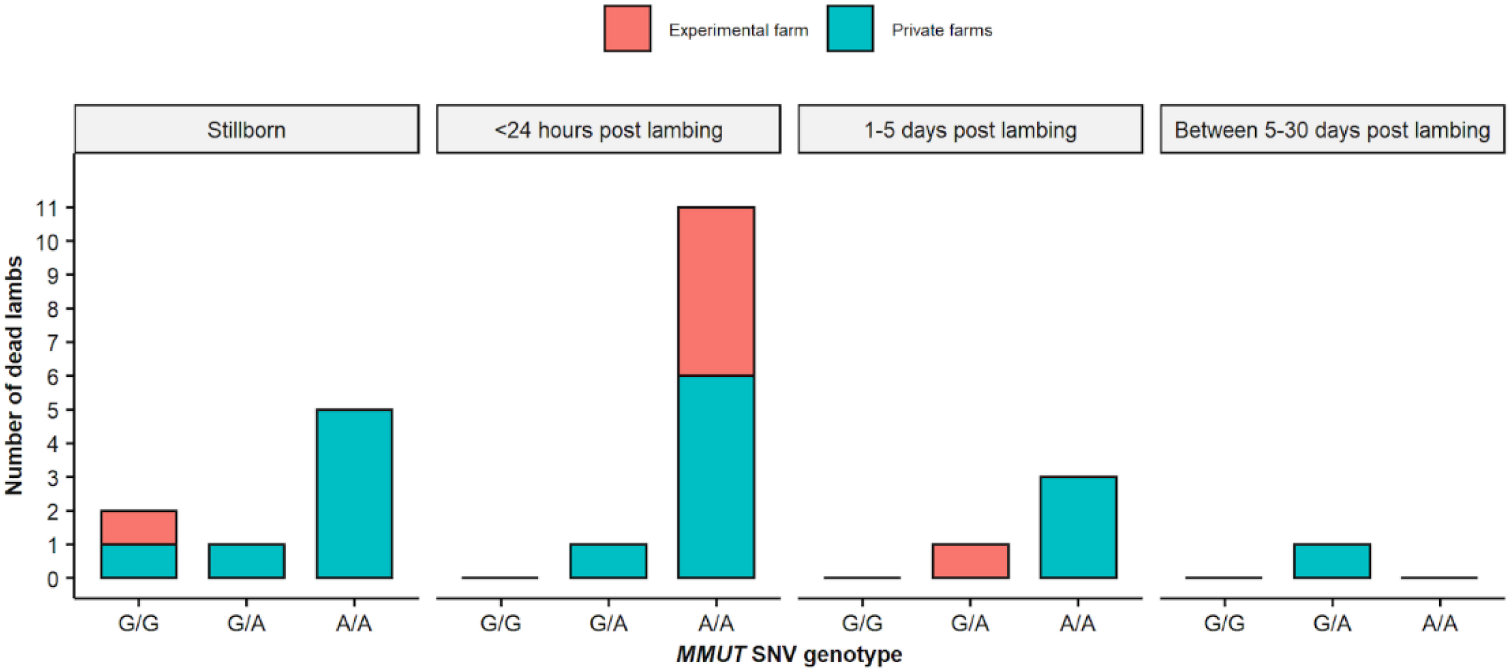
Time distribution of dead lambs in the pre-weaning period. Bat charts depending on *MMUT* SNV genotype and lambing place.

**Fig 10.**
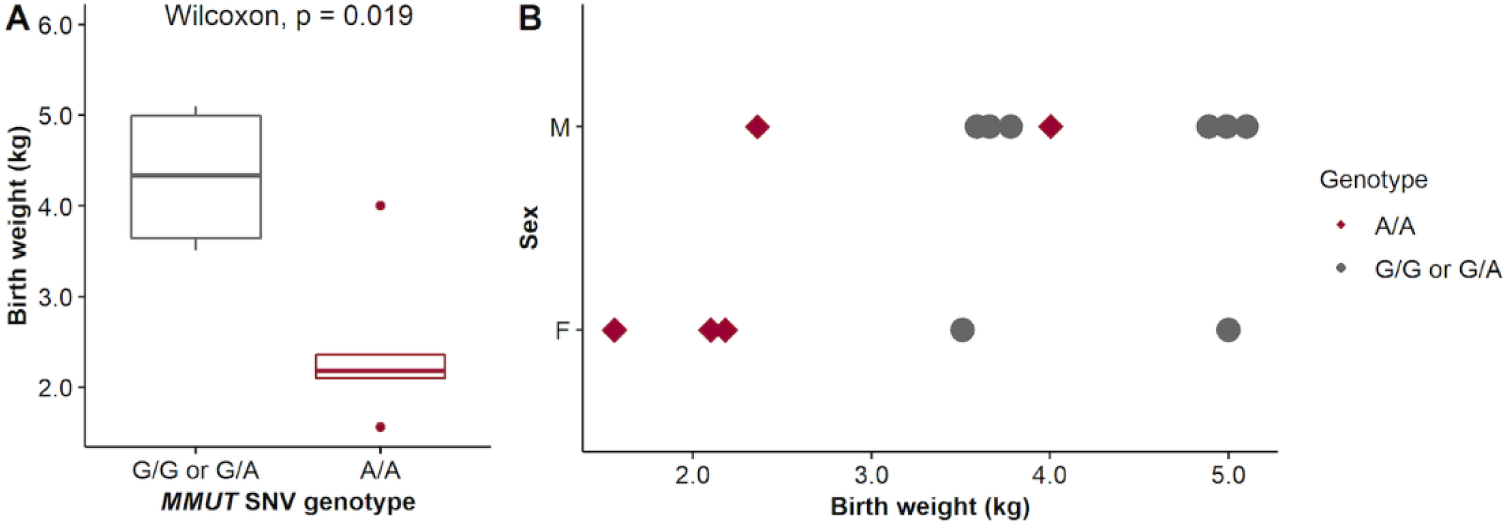
*MMUT* SNV genotype effect on lamb birth weight. (A) Boxplot representation of birth weight (n=13) according to *MMUT* genotypes (B) distribution of birth weight by sex (M: Male n=8, F: Female n=5) and genotypes. Affected homozygous lambs are in red (A/A genotype).

**Table 4.** Genotyping results of lambs generated in at-risk matings in private and experimental farms according to the sex and MMUT SNV genotype.

|  | **g.23,776,347G>A genotype in *MMUT*** |  |  |  |  |
| --- | --- | --- | --- | --- | --- |
|  | **G/G** | **G/A** | **A/A** | **-/-** | **Total** |
| <u>Experiment 1:</u><br>Private farms (n=6) |  |  |  |  |  |
| Male | 9 | 9 | 10 | 1 | 29 |
| Female | 8 | 16 | 5 |  | 29 |
| Undetermined |  |  | 1 |  | 1 |
| **Total** | **17** | **25** | **16** | **1** | **59** |
| <u>Experiment 2:</u><br>Experimental farm (n=1) |  |  |  |  |  |
| Male | 4 | 2 | 2 |  | 8 |
| Female | 0 | 2 | 3 |  | 5 |
| **Total** | **4** | **4** | **5** |  | **13** |
| <u>All</u> |  |  |  |  |  |
| Male | 13 | 11 | 12 | 1 | 37 |
| Female | 8 | 18 | 8 | 0 | 34 |
| Undetermined |  |  | 1 |  | 1 |
| **Total** | **21 (19<sup>a</sup>/2<sup>†</sup>)** | **29 (25<sup>a</sup>/4<sup>†</sup>)** | **21 (2<sup>a</sup>/19<sup>†</sup>)** | **1\* (0<sup>a</sup>/1<sup>†</sup>)** | **72** |

\*The ear punch was not available for this lamb.
<sup>a</sup> Number of alive lambs.
<sup>†</sup> Number of dead lambs.

### MMUT protein expression and activity

Based on the sheep gene atlas (http://biogps.org/sheepatlas/; accessed 17 February 2022), *MMUT* is shown to be highly expressed in kidney and liver [52]. In order to evidence a putative nonsense-mediated mRNA decay (NMD) due to the nonsense variant in *MMUT*, we evaluated the *MMUT* mRNA relative expression in kidney and liver by qPCR (Fig 11A). We evidenced a significant reduction of *MMUT* expression in liver (Wilcoxon’s non parametric test, p=0.0015) but not in kidney (Wilcoxon’s non-parametric test, p=0.66). We also assessed the MMUT protein expression from liver and kidney protein extracts collected from two homozygous A/A and two homozygous G/G dead lambs. As expected, using Western blotting, the wild type protein was expressed in kidney and liver whereas the mutated protein was not detected at least as a full-length form in both tissues (Fig 11B). We also tried to evaluate the accumulation of methylmalonic acid quantified by ELISA in urine and blood of A/A lambs collected post-mortem or soon after birth in alive animals (Fig 12) but no significant difference was observed compared to G/G or G/A lambs (Wilcoxon’s non-parametric test, p=0.54 in urine, Wilcoxon’s non-parametric test, p=0.34 in plasma).

**Fig 11.**
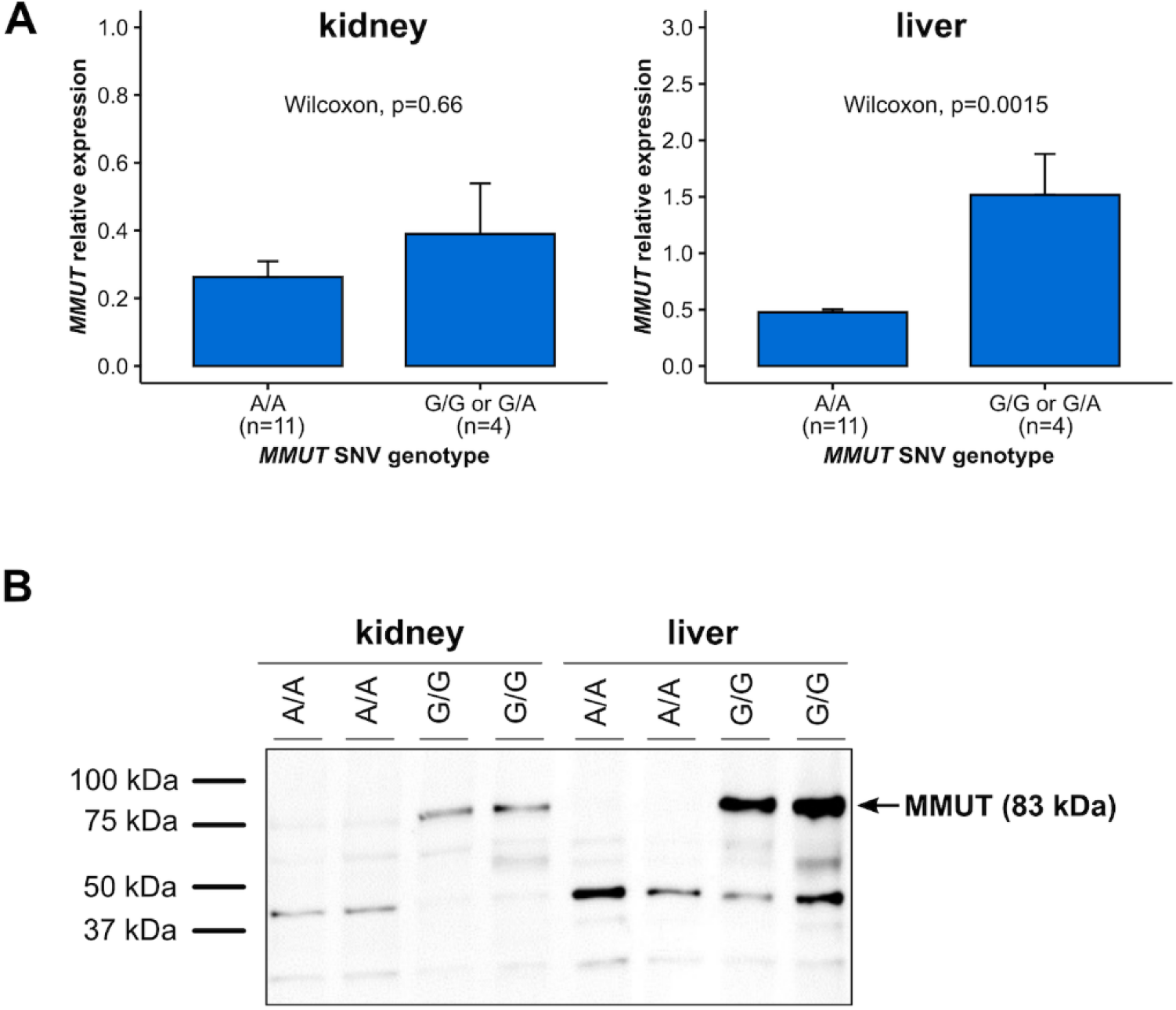
RNA and protein expression of *MMUT* in homozygous A/A lambs. *MMUT* gene expression (mean ±SEM) at mRNA (A) and protein (B) levels in kidney and liver depending on *MMUT* SNV genotypes.

**Fig 12.**
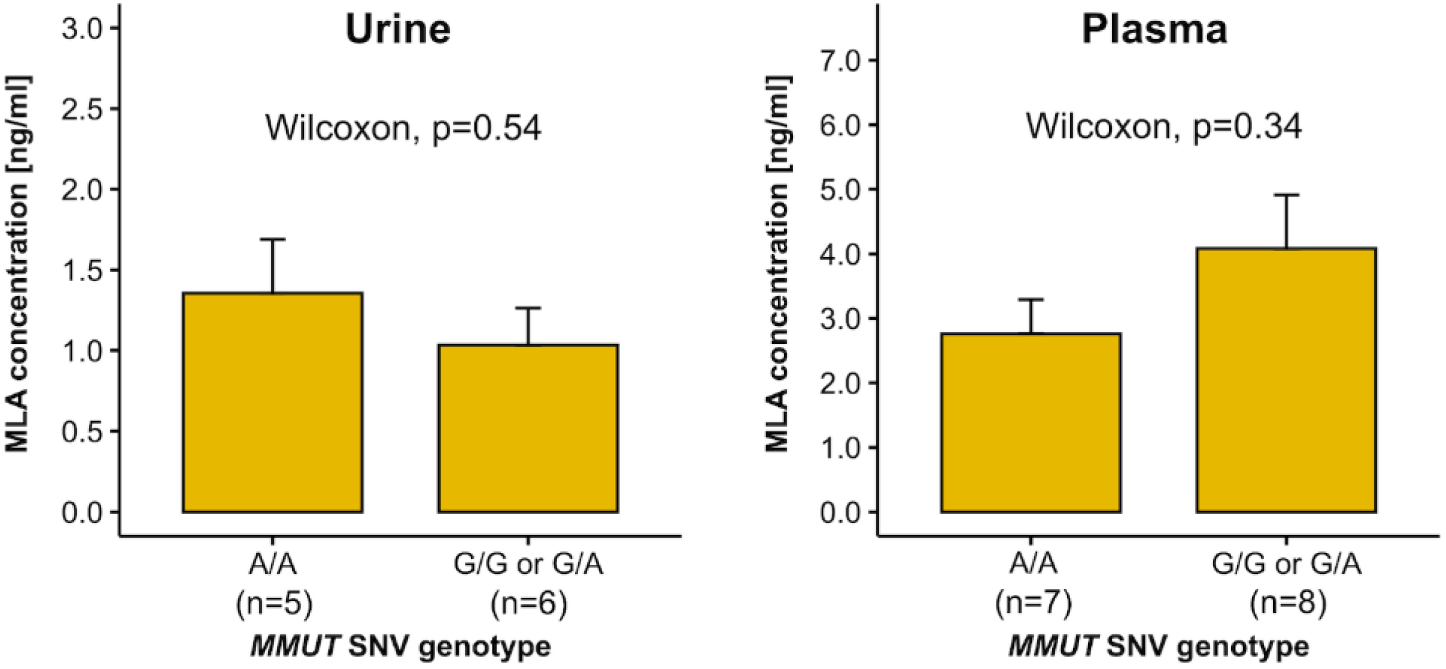
Methylmalonic acid dosage in urine and plasma of A/A lambs. Mean ± SEM of Methylmalonic acid (MLA) ELISA quantification in blood plasma and urine collected from lambs of different genotypes at the *MMUT* SNV.

## Discussion

Using a reverse genetic screen in MTR population, we successfully identified five genomic regions (named MTRDHH1 to 5) with a significant deficit of homozygous animals ranging from 84 to 100%. Compared to our previous analysis in Lacaune dairy sheep with 8 independent haplotypes, we identified less deficient haplotypes in MTR possibly due to a lower number of genotyped animals (19,102 Lacaune vs 6,845 MTR) [42]. In the MTR population, we estimated the frequencies of MTRDHH heterozygous carriers between 7.8% and 16.6%, and thus allele frequency between 3.9 and 8.3%. This is in line with an allele frequency of 5% expected from the analysis of a population of 6,000 genotyped animals [24].

We tested a putative impact of each MTRDHH on production traits to search for a selective advantage at heterozygous state. The positive effects of these DHH on selected traits were quite low and only significant for MTRDHH2 carriers on milk yield and for MTRDHH5 on fat yield. It is thus unlikely that selective advantage explains the observed DHH frequencies, or the rapid increase in MTRDHH3 frequency between 2017 and 2018. However, many examples of balancing selection for deleterious alleles have been described in livestock [53]. For example in sheep, a missense variant in *FGFR3* was associated with enhanced skeletal growth and meat yield when heterozygous while it induced chondrodysplasia (spider lamb syndrome) when homozygous (OMIA 001703-9940) [54].

The populational analysis of AIS and SBR recorded on more than 300,000 matings, allowed us to do a first sorting of the different MTRDHH based on their supposed deleterious impact on early gestation (AIS), around the time of birth (SBR), or on postnatal viability or morphological phenotypes when AIS and SBR were not altered. Accordingly, we classified the five haplotypes into three groups and we highlighted potential candidate genes according to their implication in lethal phenotypes in mouse and/or more generally involved in inherited mammalian disorders.

The first group is composed of MTRDHH3 (OAR1), MTRDHH4 (OAR10) and MTRDHH5 (OAR13) which showed no significant impact on fertility traits. We assume that these haplotypes host deleterious variants leading to postnatal lethality or morphological disorders. Within these three regions, 18 candidate genes are of interest for their implication in neonatal to juvenile lethality or associated with morphological defects counter selected at the time of candidate lamb genotyping. Candidate genes associated with postnatal lethality mainly affect metabolism (*TARS2* in MTRDHH3, *KL* in MTRDHH4) or DNA repair (*BRCA2* in MTRDHH4). Some candidate genes not necessarily lead to lethality when altered, but may affect animal welfare with the alteration of vision (*PRPF3*, *ADAMTSL4* in MTRDHH3 and *KIF3B* in MTRDHH5), neurological disorders (*PRUNE1*, *POGZ* in MTRDHH3 and *ASXL1*, *PIGU, AHCY* in MTRDHH5) or morphological/stature defects (*ITGA10*, *POLR3GL, ECM1* in MTRDHH2 and *DNMT3B*, *CEP250* in MTRDH5). However, by searching causal variants in the 22 WGS of MTR animals, we failed to detect candidate mutations in the genes listed above. In contrast, we were puzzled by the presence of *RXFP2* (MTRDHH4) and *ASIP* (MTRDHH5) as positional candidate genes already known to host variants controlling the horned/polled phenotype (OMIA 000483−9940) [48] and black coat color (OMIA 000201−9940) [49,50] in sheep, respectively. Interestingly, in MTR selection scheme, horned females and black animals do not fit with breed standard, and thus are not desirable. This may lead excluding these animals from genotyping which fully express their phenotype at the homozygous state. Accordingly, based on specific SNP markers already present on the LD SNP chip used, we were able to significantly associate MTRDHH5 with the 5pb deletion in *ASIP* leading to black coat color when homozygous. In contrast, the deficit observed for MTRDHH4 is significantly associated with the Ins allele of *RXFP2* leading to the desirable polledness trait. This intriguing observation suggests a hitchhiking phenomenon explaining the deficit [55], with the presence of a deleterious variant in linkage disequilibrium with the Ins allele of *RXFP2*.

The second group with MTRDHH2 (OAR1) associated with significant negative effects on AIS and SBR. Then, we hypothesized that the causative variant hosted by MTRDHH2 could induce embryo/fetal losses throughout the gestation period and until birth. In this region, only *SLC33A1* (*Solute Carrier Family 33 Member 1*) gene has both impact on embryonic lethality and decrease survival rate when knocked-out in mouse (MGI:1332247) and thus appears as an obvious candidate gene. In addition, variants in *SLC33A1* (OMIM 603690) are known to cause “Congenital cataracts, hearing loss, and neurodegeneration” and “Spastic paraplegia 42” phenotypes [56,57]. A study is ongoing to evidence a candidate causal variant affecting the *SLC33A1* gene in linkage disequilibrium with MTRDHH2.

MTRDHH1 (OAR20) was the only haplotype with a strong negative impact exclusively on SBR suggesting that this haplotype hosts a lethal variant affecting the perinatal period. Within the MTRDHH1 region, we identified *MMUT* (*Methylmalonyl-CoA Mutase*), *TFAP2B* (*Transcription Factor AP-2 Beta*) and *PKHD1* (*PKHD1 Ciliary IPT Domain Containing Fibrocystin/Polyductin*) as obvious candidate genes, all resulting in neonatal and/or postnatal lethality when knocked-out in mouse. *MMUT* and *PKHD1* are involved in metabolic disorders such as “Methylmalonic aciduria” (OMIM 251000) and “Polycystic kidney disease” (OMIM 263200), respectively. *TFAP2B* is involved in bone defects and heart failure (OMIM 169100). Using WGS data from 22 MTR animals and among them, two MTRDHH1 heterozygous carriers, we were able to identify four candidate polymorphisms within MTRDHH1, but only one appeared as a strong functional candidate, a SNV (G>A) located in the *MMUT* gene at position g.23,776,347 on OAR20. This SNV leads to a nonsense variation XM_004018875.4: c.1225C>T introducing a premature stop-codon (p.Gln409*). The genotyping of g.23,776,347G>A in 714 animals with a known status at MTRDHH1 indicated an almost perfect association. The only 15 discordant animals were largely explained by shorter recombinant version of the MTRDHH1 haplotype (between 2 to 31 markers surrounding the SNV). Only one MTRDHH1 heterozygous carrier did not carry the *MMUT* variant. This discrepancy could be attributed to errors from SNP array genotyping, phasing and/or imputation. Immunoblotting realized on liver and kidney proteins extracts from homozygous carrier lambs has confirmed the predicted impact of the candidate SNV on the protein. The anti-MMUT antibody has revealed a band at 83kDa as expected for the full-length MMUT polypeptide in wild-type extract. In contrast, due to the lack of the antigenic epitope (from aa 451 to 750) in the p.Gln409* truncated form, the MMUT band was not detected in homozygous carriers proving that this SNV has an effect on the functional expression of the *MMUT* gene, reinforced by a NMD phenomenon detected in liver. In order to confirm the perinatal lethality supposed for MTRDHH1 increasing SBR by 7.5%, we have managed at-risk mating between heterozygous carriers of the p.Gln409* variant. In this experiment, 84% of homozygous lambs died within the first 24 hours after birth fitting perfectly with the hypothesis and explaining the complete deficit of homozygous carriers of MTRDHH1 in the DHH analysis, due to lethality before the time of genotyping. We also evidenced that homozygous newborn lambs had a lower birth weight, an observation which could be compared to postnatal growth retardation in *Mmut* knock-out mice (MGI:5527455).

In human, numerous pathogenic variants in *MMUT* cause “Methylmalonic aciduria” (OMIM 609058, MMA, [58]), an autosomal recessive metabolism disorder. MMUT is part of a metabolic pathway starting from the degradation of amino acids (Valine, Isoleucine, Methionine and Threonine), odd-chain fatty acids, cholesterol and propionic acid to succinyl-CoA by three main enzymes: Propionyl-CoA carboxylase (PCC), Methylmalonyl-CoA epimerase (MCE) and Methylmalonyl-CoA mutase (MMUT) [59,60]. The MMUT protein is a mitochondrial enzyme that catalyze the L-methylmalonyl-CoA to succinyl-CoA, an intermediate of Krebs cycle. The isomerization of methylmalonyl-CoA requires adenosylcobalamin (AdoCbl), the cofactor form of vitamin B12 (also known as Cobalamin) [61]. In sheep, the mutant protein (p.409Gln*) does not carry the B12 binding domain (615-733 amino acids), suggesting that the AdoCbl cofactor is unable to act in the conversion of L-methylmalonyl-CoA to succinyl-CoA. When the MMUT enzyme is not functional, methylmalonic acid (MLA) accumulates in body fluids, mainly in blood and urine [60,62]. However, we failed in evidencing such MLA accumulation in plasma or urine of homozygous A/A lambs, while this was observed in homozygous knock-out mouse 24h after birth [63]. In our study, many of our urine and blood samples were collected at necropsy only after natural death of the lambs without time control, possibly affecting the results. The methylmalonic aciduria in this sheep genetic model will need further clinical investigation.

All the above elements clearly indicate that the deficit of homozygous MTRDHH1 is due to a loss-of-function mutation in the *MMUT* gene altering an essential metabolic pathway. Many reverse genetic screen approaches have also evidenced the association of DHH with mutations in genes implied in metabolism [64]. Particularly in bovine, many variants were evidenced affecting metabolic processes in several breeds such as Braunvieh (BH24/*CPT1C,* lipid metabolism) [19], Holstein (HH4/*GART*, nucleotide metabolism) [13], Montbéliarde (MH1/*PFAS,* nucleotide metabolism ; MH2/*SLC37A2,* glucose metabolism) [13,40], Normande (NH7/*CAD,* nucleotide metabolism) [26], and Simmental (SH8/*CYP2B6*, respiratory chain) [27].

Diversity analysis have also revealed the segregation of *MMUT* SNV variant in Spanish Latxa Cara Rubia (LCR) breed. French MTR and Spanish LCR are very close populations. Since 1970s, many exchanges have occurred between these population across the border, first from Spain to France during the seventies, then the reverse since the nineties [65]. The *MMUT* SNV evidenced in this study is not shared by other French sheep breeds and more largely by the individuals from the International Sheep Genome Consortium (dataset composed of 453 animals from 38 breeds all over the world, https://www.sheephapmap.org/). However, searching this dataset, we found another SNV located in ovine *MMUT* gene (rs1093255812) leading to a premature stop-gain (ENSOART00020022357.1: p.Trp43*). This variant was identified at heterozygous state in the New Zealand Coopworth breed, and as in MTR, it could have the same deleterious impact on lamb viability.

## Conclusion

In this study, we firstly identified in the MTR dairy sheep the segregation of five independent haplotypes possibly hosting five recessive deleterious alleles of which at least three have negative impact on fertility traits. Among them, we evidenced the *MMUT* c.1225T (p.409Gln*) variant associated with the MTRDHH1 haplotype causing early lamb mortality when homozygous. This could provide an excellent animal model for the study of methylmalonic aciduria occurring in human with the same recessive genetic determinism affecting *MMUT*. The MTRDHH2 and 3 represent promising haplotypes to discover other recessive lethal mutations in the near future. This reverse genetic study has also allowed us to hypothesize that two of these haplotypes, MTRDHH4 and 5, do not associate with lethal mutations but with mutations possible affecting morphological traits. The causal mutations are still to be identified since the already known mutations in *RXFP2* and *ASIP* do not fit perfectly with the segregation of these two haplotypes. Anyway, an appropriate management of these haplotypes/variants in the MTR dairy sheep selection program should increase the overall fertility and lamb survival, and may help for selection of morphological breed standards.

## Materials and methods

### Animal and genotyping data

The total dataset is composed of 6,845 genotyped Manech Tête Rousse animals (82% males and 18% females) born between 1993 and 2021 (data description in S4 Fig). Genotyping was performed at the Labogena facility (http://www.labogena.fr/) in the framework of the French dairy sheep genomic selection [66]. Both low density 15k SNP chip (SheepLD; n=2,956) and medium density 50k SNP chip (Ovine SNP50 BeadChip; n=3,889) were purchased from Illumina Inc. (San Diego, USA) (Table 5). Pedigree information was extracted from the official national database SIEOL (*Système d’Information en Elevage Ovin Laitier*, France).

**Table 5.** Description of genotyped animals.

| Year of birth | < 2017 | ≥ 2017 |  | Total |
| --- | --- | --- | --- | --- |
| Background | Research programs | Genomic selection & Research programs |  |  |
| Number of animals | 2,533 rams | 3,077 rams |  | 6,845 |
|  | 692 ewes | 543 ewes |  |  |
| SNP chip | MD | LD | MD | LD (n = 2,956) |
|  | (n=3,225) | (n = 2,956) | (n = 664) | MD (n =3,889) |
| Genotyping age | >12 months | ~15 days | 8-12 months |  |

MD: medium density (50k), LD: low density (15k)

### Genotype quality control, imputation and phasing

Quality control for each SNP was carried out following the French genomic evaluation pipeline based on (*i*) call frequency >97%, (*ii*) minor allele frequency >1%, (*iii*) respect Hardy-Weinberg equilibrium (P>10^-5^). Genotypes were phased and imputed from LD to MD using *FImpute3* [67]. The accuracy of LD to MD imputation in MTR was previously assessed and resulted in a concordance rate per animal of 98.86%, a concordance rate per SNP of 98.95% and a squared Pearson’s correlation coefficient of 93.57% between imputed and observed SNP genotypes [68]. After quality control, the 38,523 remaining autosomal SNPs were mapped onto the *Ovis aries* genome assembly Oar v3.1 (current version used in the French genomic evaluation) [69]. These SNP were also located on the genome assembly Oar_rambouillet_v1.0 (GCF_002742125.1). Genomic coordinates of both version of the sheep genome are available at https://doi.org/10.6084/m9.figshare.8424935.v2 (https://www.sheephapmap.org/).

### Detection of homozygous haplotype deficiency

Following the method used previously to detect deficit in homozygous haplotypes in sheep [42], we screened the genome of the 5,271 genotyped animals belonging to trios (77 offspring had both parents genotyped, and 4,799 offspring had both sire and maternal grandsire genotyped). Briefly, the method consists in (*i*) screening the genome using a sliding window of 20 SNP markers, (*ii*) selecting all 20 SNP haplotypes with frequency>1% on maternal phase, (*iii*) comparing the observed number *N*_Obs_(*k*) to the expected number *N*_Exp_(*k*) of homozygous offspring for each haplotype *k* using within-trios transmission probability and further considered haplotypes with P-poisson<1.9×10^-4^, (*iv*) retaining deficit between 75% and 100% defined as (*N*_*Exp*_(*k*) − *N*_*Obs*_(*k*))/*N*_*Exp*_(*k*). Finally, consecutive windows with these same parameters were clustered to define larger region called “Manech Tête Rousse Deficient Homozygous Haplotype” (MTRDHH). For each MTRDHH, the status (homozygous non-carriers, heterozygous and homozygous carriers) was thereafter determined for all 6,845 genotyped animals available. Linkage disequilibrium was estimated between two MTRDHH regions located on the same chromosome by the r2 coefficient measure as described [42].

### Analysis of fertility traits

Mating trait records of MTR between 2006 and 2019 were obtained from the national database SIEOL. Only artificial insemination success (AIS) and stillbirth rate (SBR) records from mating where both sire and maternal grand sire were genotyped (i.e. had a known status at each MTRDHH) were analyzed. AIS was coded “1” for success and “0” for failure based on lambing date according to the gestation length starting from the day of AI (151 ± 7 days; n=330,844 mating records). SBR was determined only in the AI success group, and coded “1” if there was at least one stillbirth in the litter or “0” if all lambs were born alive (n=201,637 mating records). We considered “at-risk mating” a mating between a carrier ram and a ewe from a carrier sire. We considered “safe mating” when the other combinations occurred: (i) non-carrier ram × ewe from a non-carrier sire, (ii) non-carrier ram × ewe from a carrier sire, (iii) carrier ram × ewe from a non-carrier sire. A logistic threshold binary model with a logit link function was used to compare AIS and SBR between at-risk and safe mating (lsmeans estimate), using the GLIMMIX procedure in the SAS software (version 9.4; SAS Institute Inc., Cary, NC). The fixed effects for AIS and SBR were mating type (safe or at-risk), season of AI (spring or summer), and lactation number (L1, L2, L3 and L4+). For SBR only, prolificacy of the ewe (1, 2, 3+ lambs/litter) was added as a fixed effect. For AIS and SBR, the random effect was the interaction herd×year (n=313 herds between 2006 and 2019). Traits were considered to differ significantly when the mating type fixed effect had a P-value lower than 1.0×10^-2^ after Bonferroni correction for multiple testing with a level of significance α at 5%. This threshold was obtained by dividing the level of significance α by the number of tests corresponding to the number of independent haplotypes (n=5).

### Analysis of milk parameters and total merit genomic index (ISOLg)

Daughter yield deviations (DYD) for milk traits from genotyped sires with known status at each MTRDHH were computed from official genetic evaluation (GenEval, Jouy-en-Josas, France). The DYD corresponds to the average performance of the daughters of each sire, corrected for environmental effects and the average genetic value of the dam [66]. The six traits studied were milk yield (MY), fat (FC) and protein (PC) contents, fat (FY = MY×FC) and protein (PY = MY×PC) yields, and lactation somatic cell score (LSCS) as described [42]. To compare all the traits on the same scale, each DYD was divided by its genetic standard deviation to obtain standardized DYD (sDYD). Only genotyped rams with records from at least 20 daughters were included in the analysis in order to obtain sufficiently accurate DYD values (n ∼2570 rams). Each trait was tested by variance analysis comparing MTRDHH carrier and non-carrier rams using the GLM procedure in the SAS software (version 9.4; SAS Institute Inc., Cary, NC). The fixed effects were the genetic status (carrier, non-carrier) and year of birth (2000 to 2016) to correct for annual genetic gain. Traits were considered to differ significantly when the genetic status fixed effect had a P-value lower than 1.0×10^-2^ after Bonferroni correction for multiple testing with a level of significance α at 5%. This threshold was obtained as explained above for fertility traits (n=5 independent haplotypes).

Total merit genomic index in dairy sheep (called “ISOLg”, *Index Synthétique des Ovins Laitiers*) was extracted from the official genomic evaluation (GenEval, Jouy-en-Josas, France) for all the 714 genomic candidate lambs born in 2021. ISOLg is determined by a combination of four selected traits: MY, FC, PC and LSCS. For each MTRDHH, ISOLg from heterozygous carrier and non-carrier lambs were compared with a Wilcoxon non-parametric test under the null hypothesis with a risk of α=5% using “wilcox.test” function in R software (version 4.1.3, R Core Team, 2022).

### Identification of positional and functional candidate genes

All annotated genes located in the MTRDHH region extended by 1 Mb from each side were extracted from the ovine genome Oar_rambouillet_v1.0 (OAR1: NC_040252.1, OAR10: NC_040261.1, OAR13: NC_040264.1 and OAR20: NC_040271.1) using CLC export annotation function (QIAGEN CLC Main Workbench 7.9). Genes with a known knock-out phenotype in mouse including mortality and aging (embryonic, prenatal, perinatal, neonatal, postnatal, preweaning, premature death and decreased survival rate) or associated with mammalian genetic disorders were sorted using “biomaRt” R package (version 2.52.0, https://doi.org/doi:10.18129/B9.bioc.biomaRt) extracted from Mouse Genome Informatics (MGI, http://www.informatics.jax.org), International Mouse Phenotyping Consortium (IMPC, https://www.mousephenotype.org), Online Mendelian Inheritance in Man (OMIM, https://omim.org) and Online Mendelian Inheritance in Animal (OMIA, https://omia.org) databases (last accession on 27 May 2022). Relevant candidate genes were presented as a heatmap using “pheatmap” R package (version 1.0.12).

### Whole genome sequencing data

Publicly available data of 100 ovine short-read Illumina HiSeq/NovaSeq whole genome sequences (WGS) from 14 breeds generated in various INRAE and Teagasc research projects were used for variant calling. Among them, 22 WGS were obtained from MTR dairy sheep also genotyped with the MD SNP chip. A description of the different breeds and the accession numbers of sequencing raw data are available in S3 Table.

### WGS variant calling and filtering

Reads mapping, variant calling and functional annotation were performed using Nextflow v20.10.0 and Sarek v2.6.1 pipelines for the 100 short-read WGS as previously described [43]. Regions of interest were extracted using SnpSift Filter, part of the SnpEff toolbox [51]. Candidate variants were filtered based on the correlation between haplotype status (homozygous non-carriers, heterozygous and homozygous carriers encoded as 0, 1 and 2, respectively) and allele dosage for bi-allelic variants (also encoded 0, 1 and 2) using geno--r2 command of VCFtools [70].

### Specific variant genotyping assays

Genotyping at the *RXFP2* (1.8 kb InDel) and *ASIP* (5pb InDel) loci were directly obtained from the LD SNP chip based on the specific markers described in S4 Table. The NC_040264.1:g.66,474,980T>A in the last exon of *ASIP* and the SNP NC_040271.1: g.23,776,347G>A in *MMUT* were both genotyped by PACE (PCR allele competitive extension) analysis. Fluorescent PACE analysis was done with 15 ng of purified DNA using the PACE-IR 2x Genotyping Master mix (3CR Bioscience) in the presence of 12 µM of a mix of extended allele specific forward primers and 30 µM of common reverse primers in a final volume of 10 μL (primer sequences described in S5 Table). The touch-down PCR amplification condition was 15 min at 94°C for the hot-start activation, 10 cycles of 20 s at 94°C, 54–62°C for 60 s (dropping 0.8°C per cycle), then 36 cycles of 20 s at 94°C and 60 s at 54°C performed on an ABI9700 thermocycler followed by a final point read of the fluorescence on an ABI QuantStudio 6 real-time PCR system and using the QuantStudio software 1.3 (Applied Biosystems).

The presence of the *RXFP2*, *ASIP* and *MMUT* variants was checked in a DNA set of the 2021 cohort of 714 MTR male lamb candidates for genomic selection. DNA was extracted by Labogena (Jouy-en-Josas, France) on behalf of the MTR breed industry. A DNA diversity panel of 851 animals from 25 French sheep breeds [71] and 3 Spanish sheep breeds was also specifically genotyped for the *MMUT* variant.

### Generation of homozygous lambs

Blood samples (3 ml) were first collected from 181 ewes, daughters of MTRDHH1 carrier sires, located in 6 private farms by jugular vein puncture with the Venoject system containing EDTA (Terumo, Tokyo, Japan) and directly stored at -20°C. Among them, 82 ewes were genotyped as heterozygous carriers of the *MMUT* variant and were selected to be inseminated with *MMUT* variant heterozygous rams (at-risk matings). Experiment 1 (n=73 ewes) was performed in 6 private farms in Pays-Basque (France) and Experiment 2 (n=9 ewes) was performed at the INRAE experimental farm of Langlade under the agreement number E31429001 (Pompertuzat, France). Experimental design is described in Fig 13. An ultrasound diagnosis of gestation was realized between 45 and 60 days after AI. Gestations were followed and each lamb was monitored form birth to weaning. Ear biopsies (1 mm^3^) from the 72 lambs born in both experiments were obtained with a tissue sampling unit (TSU, Allflex Europe, Vitré, France) and directly placed in the TSU storage buffer at 4°C. Ear biopsies were placed twice consecutively in 180 μL of 50 mM NaOH, heated 10 min at 95°C, neutralized with 20 μL of 1 M Tris-HCl, and then vortexed during 10s. All neutralized samples were used for direct genotyping without DNA purification as described [72]. In Experiment 2, all lambs were weighted at birth. Biological samples (plasma, urine, liver and kidney) were collected on animals as described in Fig 13 and frozen at -80°C until use.

**Fig 13.**
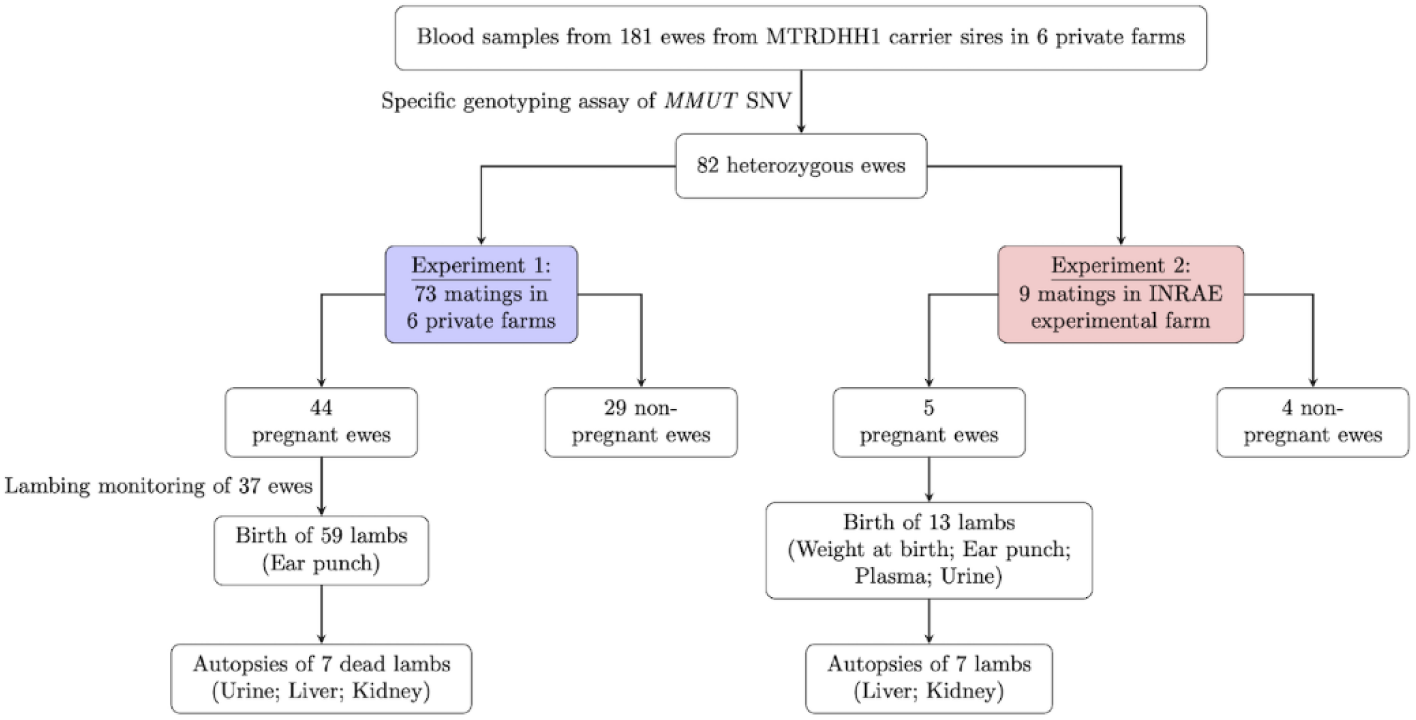
Experimental design to generate *MMUT* homozygous variant lambs.

### Immunoblot analysis

Frozen kidney and liver tissues were crushed in liquid nitrogen using a Mixer Mill during 30 seconds at 30 Hz (MM400, Retsch technology), and 12 mg of tissue powder were lysed with 500µL of RIPA solution (Ref#R0278, Sigma-Aldrich). The protein extracts were centrifuged for 20 minutes at 16 000g and 4°C, and protein concentration in the supernatant was determined using BCA protein assay Kit (Ref#K812-1000, Biovision). Each protein sample (45 µg) was denatured and reduced in Laemmli buffer (62.5mM TRIS pH 6.8, 2% SDS; 10% glycerol) containing 5% ß-mercaptoethanol before SDS-PAGE on a 4-15% polyacrylamide gel (Bio-Rad). Proteins were transferred onto a nitrocellulose membrane blocked with AdvanBlock-Chemi blocking solution (Advansta) during 1 hour at room temperature. After washing in PBS-0.1% Tween 20, the membrane was incubated overnight à 4°C with a rabbit polyclonal anti-MMUT primary antibody (MUT Rabbit pAb, Ref#A3969, ABclonal), followed (after washing) by 1h with a rabbit polyclonal anti-Actin primary antibody (Ref#A2066, Sigma-Aldrich) both at 1/1000 in blocking solution. Revelation of the primary antibodies was performed by incubation with goat anti-rabbit Horseradish peroxidase-conjugated secondary antibody (Ref#A0545, Sigma-Aldrick) at 1/10000 ratio in blocking solution for 1 hour at room temperature, followed by enhanced chemiluminescence detection (WesternBright Quantum HRP substrate, Advansta) on a ChemiDoc Touch low-light camera (Bio-Rad) in automatic mode.

### RNA extraction, reverse transcription and quantitative PCR

Total RNA was extracted from 80 mg of frozen kidney (n=15) and liver tissue powders (n=15) in 1mL Trizol reagent (Invitrogen, #Ref 15596-018) and isolated from Nucleospin® RNA II kit (Macherey-Nagel, #Ref 740955.50) according to the manufacturer’s protocol and including DNAseI digestion treatment. The RNAs were quantified by spectrophotometry (NanoDrop® ND-8000 spectrophotometer, ThermoFischer) and stored at -80°C. Reverse transcription was carried out from 1µg of total RNA in solution with anchored oligo(dT) T22V (1µL at 10μM), random oligo-dN9 (1µL at 10μM) and dNTPs (2µL at 10 mM) in a reaction volume of 10 μL. This mixture was incubated at 65°C for 5 min in an ABI2700 thermocycler (Applied Biosystems) then ramped down to 4°C. A second reaction mixture (8 μL/reaction) containing the reaction buffer (5µL of First strand Buffer 5X, Invitrogen, France), DTT (Dithiothreitol, 1µL at 0.1M), Rnasine (1µL, 40 units/µL, Promega, France) and Superscript II reverse transcriptase (1µL, 200 units/µL, Invitrogen, France) was added to the denatured RNA solution (final volume reaction of 18μL) then incubated for 50 minutes at 42°C and placed for 15 minutes at 70°C. The complementary DNA (cDNA) solution obtained was directly diluted at 1:5 ratio and stored at -20°C. For each pair of primers, amplification efficiency was evaluated by *E* = *e*^−1/*α*^ which α is the slope of a linear curve obtained from cDNA serial dilution (1:5 to 1:80) and corresponding Ct (cycle threshold) values. Quantitative PCR (qPCR) was performed using 3µL of cDNA at 1:20 ratio, 5µL of SYBR Green real-time PCR Master Mix 2X (Applied Biosystems) and 2µL of primers at 3µM in a total reaction volume of 10µL on qPCR was realized on a QuantStudio 6 Flex Real-Time PCR system (ThermoFisher). Each sample was tested in duplicate. RNA transcript abundance was quantified using the ΔΔ*Ct* method corrected by four reference genes (*GAPDH*, *YWHAZ, RPL19* and *SDHA*) and a calibrator sample. Primers were design using Beacon Designer™ 8 (Premier Biosoft). The list of qPCR primer sequences, amplification length and amplification efficiency used is available in S5 Table.

### Methylmalonic acid dosage assay

Dosage of methylmalonic acid (MLA) was performed on 11 urine and 15 plasma samples collected on lambs. MLA dosage was performed using Sheep Methylmalonic Acid Elisa kit (MyBioSource, Ref# MBS7266308) following the manufacture’s protocol starting with 100µL of samples. The incubation phase with specific antibody was performed overnight at 4°C. The optical density at 450 nm was determined on a GloMax®-Multi Detection System (Promega).

### Ethic statement

The experimental procedures on animals were approved (approval numbers APAFIS#30615-2021032318054889 v5) by the French Ministry of Teaching and Scientific Research and local ethical committee C2EA-115 (Science and Animal Health) in accordance with the European Union Directive 2010/63/EU on the protection of animals used for scientific purposes.

## Acknowledgments

We are grateful to the genotoul bioinformatics platform Toulouse Occitanie (Bioinfo Genotoul, https://doi.org/10.15454/1.5572369328961167E12) for providing help and/or computing and/or storage resources. The authors acknowledge the breeding confederations CDEO (Centre Départemental de l’Elevage Ovin) and CONFELAC (Confederación de Asociaciones de Criadores de Ovino de Razas Latxa y Carranzana) for providing access to private genomic data and/or biological samples. We thank also the breeders involved in the project and Soline Szymczak for help with genotyping.

## Funding

This research has received funding from the European Union’s Horizon 2020 research and innovation program under the Grant Agreement No. 772787 (SMARTER) and PRESAGE project (CASDAR n°20ART1532777, the responsibility of the French ministry of Agriculture and Food cannot be engaged). MB was supported by a Ph.D. grant for the HOMLET program co-funded by APIS-GENE and Région Occitanie.

## Supporting information captions

**S1A Table. Clustering HHD into MTRDHH regions.** Table shows all significant haplotypes of 20 markers (150 HHD with frequency > 1%, P-value < 1.9 × 10^−4^ and deficit ≥ 75%). As described in the Materials and methods section, the 150 HHD could be clustered into 5 MTRDHH regions. **S1B Table. SNPs defining the MTRDHH regions.** Table gives the position of each SNP within MTRDHH regions according to the sheep reference genomes Oar_v3.1, Oar_rambouillet_v1.0 and ARS-UI_Ramb_v2.0, and the phased alleles of each deficient haplotype.

**S1 Fig. Total merit genomic index (ISOLg) for the 2021 cohort genomic lambs (n=714).** (A) Distribution of ISOLg in MTR dairy sheep. The ISOLg is determined by a combination of four selected traits: MY, FC, PC and LSCS. Comparison of ISOLg according to each DHH status, (B) MTRDHH1, (C) MTRDHH2, (D) MTRDHH3, (E) MTRDHH4, (F) MTRDHH5.

**S2 Table. List of the 408 protein coding genes located in the five MTRDHH extended by 1 Mb on each side.** Information on mouse phenotypes and association with mammalian disorders were extracted for each gene using several databases: MGI: www.informatics.jax.org; IMPC: https://www.mousephenotype.org); OMIM: Online Mendelian Inheritance in Man (https://omim.org) and OMIA: Online Mendelian Inheritance in Animal (https://omia.org). For mouse phenotypes associated with lethality, these were separated by developmental stages and encoded 0 (no lethality) or 1 if the gene is involved in lethality at the given stage.

**S2 Fig. Association between MTRDHH4-5 and causal variants involved in morphological traits from the 22 sequenced MTR animals.** (A) Association between MTRDHH4 status and 1.8kb insertion in the 3’-UTR of *RXFP2* associated with polledness. Association between MTRDHH5 status and (B) OAR13:g.66,475,132_66,475,136del and (C) OAR13:g.66,474,980T>A in *ASIP* associated with black coat color. Coordinates refer to sheep genome Oar_rambouillet_v1.0.

**S3 Fig. MTRDHH1 recombinant haplotypes from 15 animals showing mismatch between MTRDHH1 status and *MMUT* SNV genotype.** MTRDHH1/+ and +/+ refer to heterozygous and non-carriers of MTRDHH1, respectively. The grey column represents the localization of the *MMUT* SNV (g.23,776,347G>A) within the MTRDHH1 haplotype. For each animal, only the phase supposed to host the *MMUT* SNV variant allele is represented. The blue color indicates the portion of local haplotype matching with the MTRDHH1 haplotype.

**S4 Fig. Distribution of genotyped animals over time.** The bar charts represent the number of genotyped animals according to sex and year of birth. The genomic selection in MTR dairy sheep was implemented in 2017.

**S3 Table. EMBL-EBI accession numbers of the 100 whole-genome sequences used in the analysis.**

**S4 Table. Markers on SheepLD chip for genotyping at the *RXFP2* (1.8 kb InDel) and *ASIP* (5pb InDel) loci.**

**S5 Table. List of PCR primer sequences.**

## Notes

### Competing Interest Statement

The authors have declared no competing interest.

### Summary of Updates

Minor text edition and addition of an author.

