## Supplementary figures and images for "Homozygous haplotype deficiency in Manech Tête Rousse dairy sheep revealed a nonsense variant in *MMUT* gene affecting newborn lamb viability"

### Supplemental Figure 1

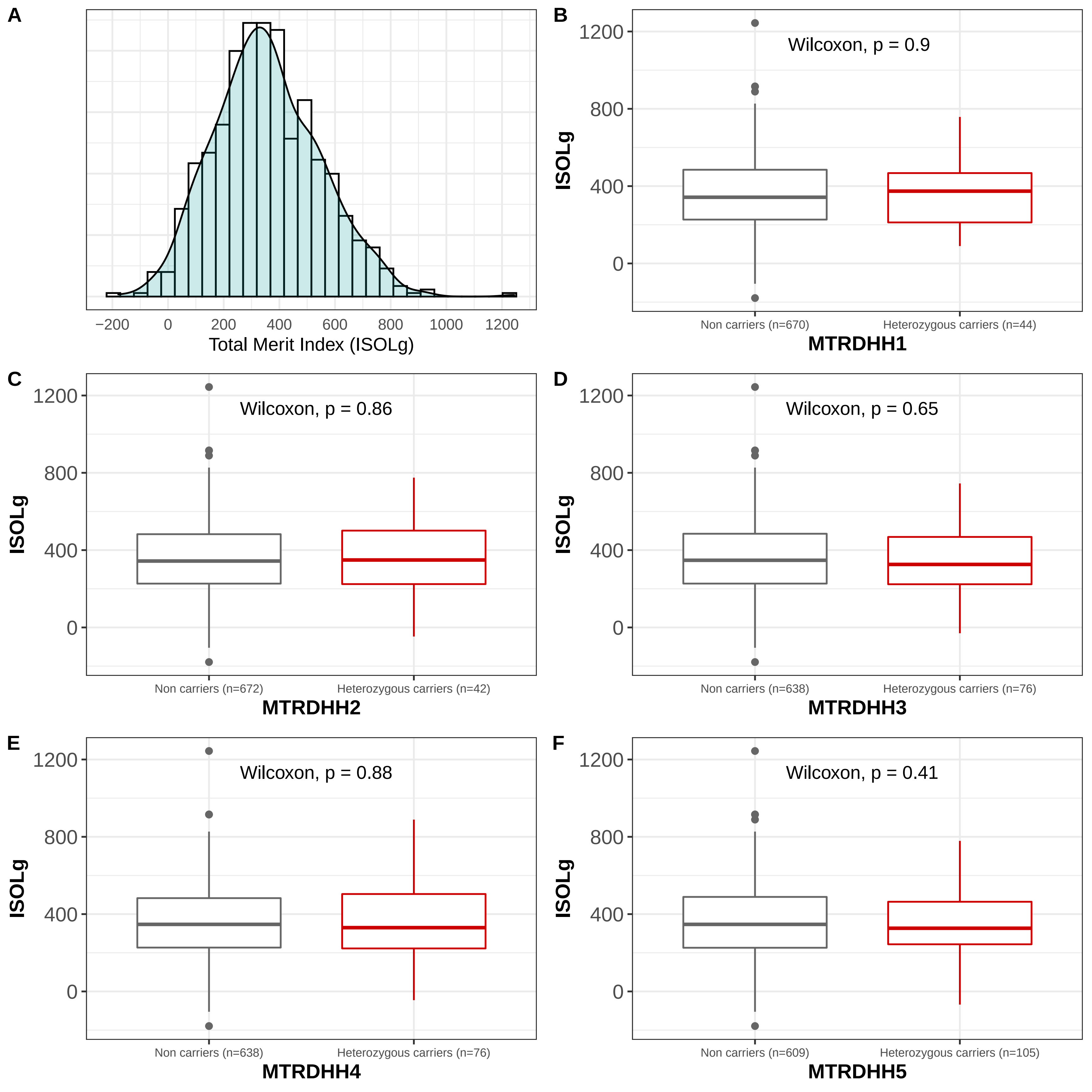

### Supplemental Figure 2

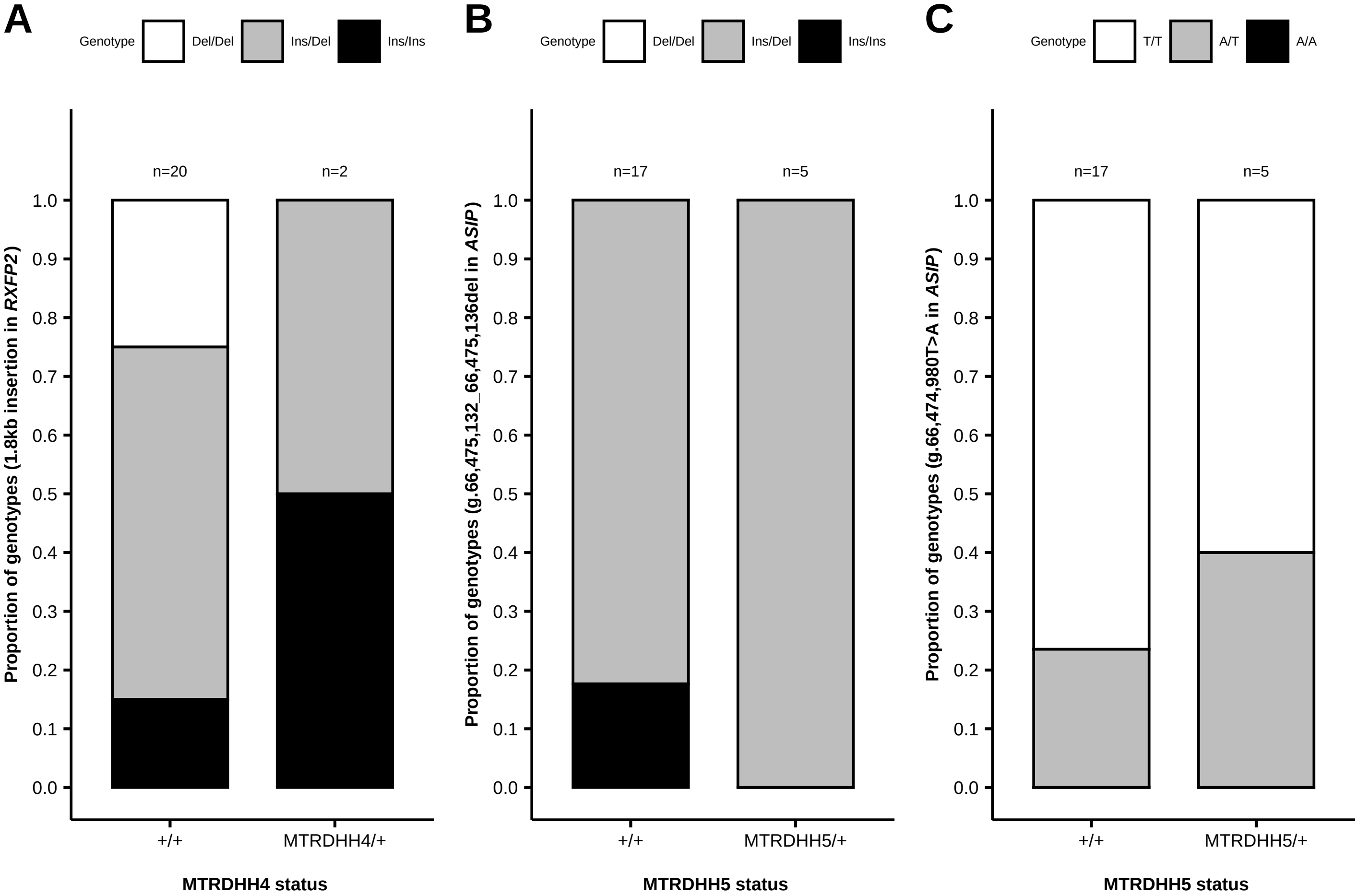

### Supplemental Figure 4

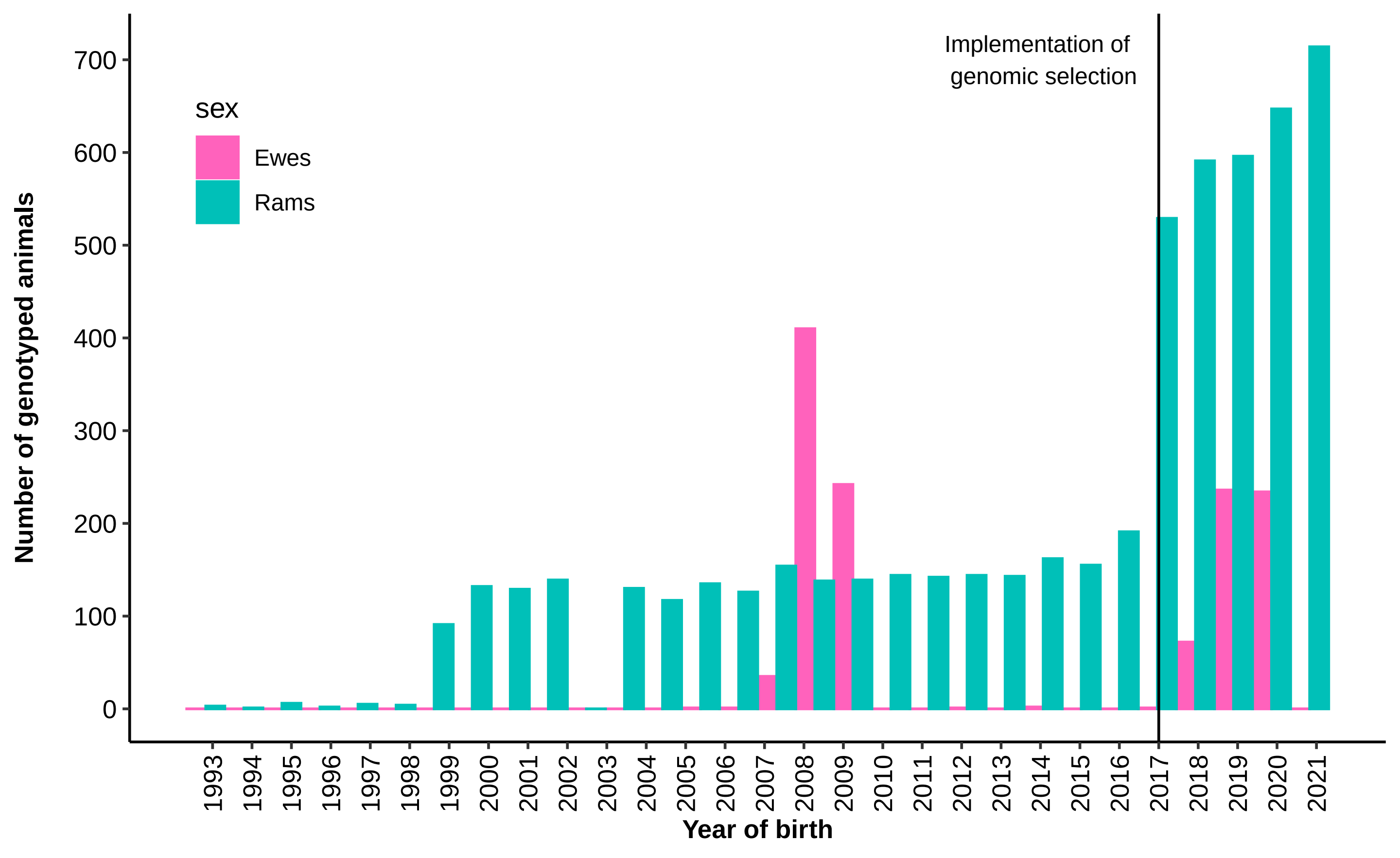
