## Supplemental Figure 3 for "Homozygous haplotype deficiency in Manech Tête Rousse dairy sheep revealed a nonsense variant in *MMUT* gene affecting newborn lamb viability"

| **MTRDHH1 haplotype** (32 markers) | | | SNP1 | SNP2 | SNP3 | SNP4 | SNP5 | SNP6 | SNP7 | SNP8 | SNP9 | SNP10 | SNP11 | SNP12 | SNP13 | | SNP14 | SNP15 | SNP16 | SNP17 | SNP18 | SNP19 | SNP20 | SNP21 | SNP22 | SNP23 | SNP24 | SNP25 | SNP26 | SNP27 | SNP28 | SNP29 | SNP30 | SNP31 | SNP32 |
|---|---|---|---|---|---|---|---|---|---|---|---|---|---|---|---|---|---|---|---|---|---|---|---|---|---|---|---|---|---|---|---|---|---|---|---|
| | | | **G** | **G** | **G** | **A** | **G** | **G** | **G** | **G** | **A** | **A** | **A** | **G** | **G** | | **G** | **A** | **A** | **G** | **A** | **A** | **G** | **C** | **A** | **G** | **G** | **A** | **G** | **C** | **G** | **A** | **G** | **G** | **A** |
| Animal | MTRDHH1 status | *MMUT* genotype | s75212.1 [A/G] | OAR20_22661712_X.1 [A/G] | OAR20_22784700.1 [A/G] | OAR20_22851804.1 [A/C] | s39376.1 [G/A] | OAR20_22932011_X.1 [A/G] | OAR20_22996638.1 [G/A] | OAR20_23028179.1 [G/A] | OAR20_23063796.1 [A/C] | OAR20_23099257_X.1 [G/A] | OAR20_23116747.1 [A/G] | s66452.1 [A/G] | OAR20_23240800.1 [A/G] | ***MMUT* SNV (G>A)** | s04959.1 [G/A] | s12178.1 [A/G] | OAR20_23408992.1 [A/C] | OAR20_23455049.1 [A/G] | OAR20_23481589_X.1 [G/A] | OAR20_23552034_X.1 [A/G] | OAR20_23606688.1 [A/G] | OAR20_23662826.1 [A/C] | OAR20_23756997.1 [A/G] | OAR20_23854908.1 [G/A] | OAR20_23878963.1 [A/G] | OAR20_23898925.1 [A/G] | OAR20_24029500.1 [G/A] | OAR20_24071930.1 [A/C] | OAR20_24089242_X.1 [G/A] | OAR20_24307087.1 [G/A] | OAR20_24357620.1 [A/G] | OAR20_24553232.1 [G/A] | OAR20_24583511.1 [A/G] |
| Animal 1 | +/+ | G/A | | | | | | | | | | | | | | A | | | | | | | | | | | | | | | | | | | |
| Animal 2 | +/+ | G/A | | | | | | | | | | | | | | A | | | | | | | | | | | | | | | | | | | |
| Animal 3 | +/+ | G/A | | | | | | | | | | | | | | A | | | | | | | | | | | | | | | | | | | |
| Animal 4 | +/+ | G/A | | | | | | | | | | | | | | A | | | | | | | | | | | | | | | | | | | |
| Animal 5 | +/+ | G/A | | | | | | | | | | | | | | A | | | | | | | | | | | | | | | | | | | |
| Animal 6 | +/+ | G/A | | | | | | | | | | | | | | A | | | | | | | | | | | | | | | | | | | |
| Animal 7 | +/+ | G/A | | | | | | | | | | | | | | A | | | | | | | | | | | | | | | | | | | |
| Animal 8 | +/+ | G/A | | | | | | | | | | | | | | A | | | | | | | | | | | | | | | | | | | |
| Animal 9 | +/+ | G/A | | | | | | | | | | | | | | A | | | | | | | | | | | | | | | | | | | |
| Animal 10 | +/+ | G/A | | | | | | | | | | | | | | A | | | | | | | | | | | | | | | | | | | |
| Animal 11 | +/+ | G/A | | | | | | | | | | | | | | A | | | | | | | | | | | | | | | | | | | |
| Animal 12 | +/+ | G/A | | | | | | | | | | | | | | A | | | | | | | | | | | | | | | | | | | |
| Animal 13 | +/+ | G/A | | | | | | | | | | | | | | A | | | | | | | | | | | | | | | | | | | |
| Animal 14 | +/+ | G/A | | | | | | | | | | | | | | A | | | | | | | | | | | | | | | | | | | |
| Animal 15 | MTRDHH1/+ | G/G | | | | | | | | | | | | | | G | | | | | | | | | | | | | | | | | | | |
