## Supplemental Table 3 for "Homozygous haplotype deficiency in Manech Tête Rousse dairy sheep revealed a nonsense variant in *MMUT* gene affecting newborn lamb viability"

**S3 Table.** EMBL-EBI accession numbers of the 100 whole-genome sequences used in the analysis

| Breed | Number of animals | ENA run accession | EBI project accession | Associated references |
|---|---|---|---|---|
| Belclare | 2 | SRR14934360; SRR14935129 | PRJNA698548 | |
| Berrichon du Cher | 3 | ERS1205899; ERS1205900; ERS1205901 | PRJEB14418 | [1] |
| Cambridge | 7 | ERR1419201; ERR1419202; ERR1419203; ERR1419204; ERR1419205; ERR1419206; ERR1419207 | PRJEB14098 | |
| Charollais | 1 | SRR14934359 | PRJNA698548 | |
| Lacaune (Dairy) | 31 | ERR3276357; ERR3276358; ERR3276359; ERR3276360; ERR3276361; ERR3276362; ERR3276363; ERR3276364; ERR3276365; ERR3276366; ERR3276367; ERR3276368; ERR3276369; ERR3276370; ERR3276371; ERR3276372; ERR3276373; ERR3276374; ERR3276375; ERR3276376; ERR3276377; ERR3276378; ERR3276379; ERR7891349; ERR7891350; ERR7891351<br>ERR968423; ERR968424; ERR968425<br>SRR501850; SRR501851 | PRJEB32110<br>PRJEB9911<br>PRJNA160933 | [2,3] |
| Lacaune (Meat) | 3 | ERR3276380; ERR3276381; ERR3276382 | PRJEB32110 | |
| Manech Tête Rousse | 22 | ERR3712282; ERR3712283; ERR3712284; ERR3712285*; ERR3712286; ERR3712287; ERR3712288; ERR3712289; ERR3712290; ERR3712291; ERR3712292; ERR3712293; ERR3712294; ERR3712295; ERR3712296; ERR3712297; ERR3712298; ERR3712299; ERR3712300; ERR3712301*; ERR7889920; ERR7889921 | PRJEB35682 | |
| Martinik Blackbelly | 1 | ERR3255914 | PRJEB31930 | |
| Noire du velay | 2 | ERR3828659; ERR3828660 | PRJEB35553 | [4] |
| Romane | 4 | ERS1205902; ERS1205903; ERR2818429; ERR2818430 | PRJEB14418 | [1] |
| Romane x Martinik Blackbelly | 13 | ERR3255915; ERR3255916; ERR3255917; ERR3255918; ERR3255919; ERR3988554; ERR3988555; ERR3988556; ERR3988557; ERR3988558; ERR3988559; ERR3988560; ERR3988561 | PRJEB31930 | |
| Romanov | 2 | ERS1205904; ERS1205905 | PRJEB14418 | [1] |
| Suffolk | 2 | SRR14934357; SRR14934358 | PRJNA698548 | |
| Texel | 2 | SRR14934355; SRR14934356 | PRJNA698548 | |
| Vendéen | 5 | ERR4236129; ERR4236130; ERR4236131<br>SRR14934353; SRR14934354 | PRJEB37460<br>PRJNA698548 | [5] |
| **Total** | **100** | | | |

* MTRDHH1 heterozygous carrier
