## Supplemental Table 4 for "Homozygous haplotype deficiency in Manech Tête Rousse dairy sheep revealed a nonsense variant in *MMUT* gene affecting newborn lamb viability"

**S4 Table. Markers on SheepLD chip for genotyping at the *RXFP2* (1.8 kb InDel) and *ASIP* (5pb InDel) loci.**

| Mutation | Markers (LD chip) | Possible genotype at the mutation |
|---|---|---|
| OAR13:g.66,475,132_66,475,136del | oar3_OAR13_63047423 (D/I) | Ins/Ins<br>Ins/Del<br>Del/Del |
| OAR10: 1.8kb insertion in the 3'-UTR of *RXFP2* | RXFP2_insert_L1 (A/G)<br>RXFP2_insert_R1 (A/T) | Ins/Ins (RXFP2_insert_L1 [G/G] & RXFP2_insert_R1 [A/A])<br>Ins/Del (RXFP2_insert_L1 [A/G] & RXFP2_insert_R1 [A/T])<br>Del/Del (RXFP2_insert_L1 [A/A] & RXFP2_insert_R1 [T/T]) |

The interpretation of the 1.8kb insertion in *RXFP2* is possible by the combination of two markers RXFP2_insert_L1 and RXFP2_insert_R1.
