## Supplemental Table 5 for "Homozygous haplotype deficiency in Manech Tête Rousse dairy sheep revealed a nonsense variant in *MMUT* gene affecting newborn lamb viability"

### S5 Table. List of PCR primer sequences

| Application | Gene symbol | Sequence (5’→3’) | fragment size (pb) | Efficiency |
|---|---|---|---|---|
| Terra PCR | *MMUT*<br>NC_040271.1 | **F:** GCAGATTTTGGCAAAGAAAG<br>**R:** ACCTTCAAGGCAGCATCATA | 402 | - |
| PACE PCR | *MMUT*<br>NC_040271.1 | **F1:** GAAGGTGACCAAGTTCATGCTAATCCCAGATTCTTCTTGAATAATGATTTG<br>**F2:** GAAGGTCGGAGTCAACGGATTGAATCCCAGATTCTTCTTGAATAATGATTTA<br>**R:** GTGAAAAGTGCTCGAATTGCCAGGAA | 81/82 | - |
| | *ASIP (SNP T>A)*<br>NC_040264.1 | **F1:** GAAGGTGACCAAGTTCATGCTTCCGCAGCGCCTGCTCCT<br>**F2:** GAAGGTCGGAGTCAACGGATTCCGCAGCGCCTGCTCCA<br>**R:** CGCGCTCAGCAGGTGGGGTT | 70 | - |
| Quantitative PCR | *MMUT*<br>*XM_004018875* | **F:** GCTCATCAAGGAACTCAA<br>**R:** TTCAATATCATCAAGCACTTG | 169 | 1.80 |
| | *GAPDH*<br>NM_001190390 | **F:** CGACTTCAACAGCGACACTC<br>**R:** TGCTGTAGCCGAATTCATTG | 113 | 1.90 |
| | *YWHAZ*<br>NM_001267887 | **F:** ATTAAGTGAAGAGTCATACAA<br>**R:** GTATCCGATGTCCACAAT | 81 | 2.00 |
| | *RPL19*<br>XM_004012836 | **F:** AATGCCAATGCCAACTC<br>**R:** CCCTTTCGCTACCTATACC | 149 | 2.00 |
| | *SDHA*<br>XM_027980212 | **F:** GAATGGTCTGGAACACTG<br>**R:** AGTAATCGTACTCGTCAAC | 156 | 2.00 |

**F:** Forward primer, **R:** Reverse primer.
